# Dynamic functional connectivity correlates of trait mindfulness in early adolescence

**DOI:** 10.1101/2024.07.01.601544

**Authors:** Isaac N. Treves, Hilary A. Marusak, Alexandra Decker, Aaron Kucyi, Nicholas A. Hubbard, Clemens C.C. Bauer, Julia Leonard, Hannah Grotzinger, Melissa A. Giebler, Yesi Camacho Torres, Andrea Imhof, Rachel Romeo, Vince D. Calhoun, John D.E. Gabrieli

**Affiliations:** McGovern Institute for Brain Research, Massachusetts Institute of Technology, Cambridge, MA; Department of Brain and Cognitive Sciences, Massachusetts Institute of Technology, Cambridge, MA; Department of Psychiatry and Behavioral Neurosciences, Wayne State University School of Medicine, Detroit, MI; Department of Psychological & Brain Sciences, Drexel University, Philadelphia, PA; Department of Psychology, University of Nebraska, Lincoln, NE; Department of Psychology, Northeastern University, Boston, MA; Department of Psychology, Yale University, New Haven, CT; Department of Psychological & Brain Sciences, University of California, Santa Barbara, CA; Teachers College, Columbia University, New York, NY; Department of Psychology, University of Oregon, Eugene, OR; Departments of Human Development & Quantitative Methodology and Hearing & Speech Sciences, and Program in Neuroscience & Cognitive Science, University of Maryland College Park, Baltimore, MD; Tri-institutional Center for Translational Research in Neuroimaging and Data Science (TReNDS), Georgia State, Georgia Tech, and Emory, Atlanta, GA

**Keywords:** Brain states, dynamics, hyperconnectivity, mindfulness, adolescents, trait mindfulness

## Abstract

**Background:** Trait mindfulness, the tendency to attend to present-moment experiences without judgement, is negatively correlated with adolescent anxiety and depression. Understanding the neural mechanisms underlying trait mindfulness may inform the neural basis of psychiatric disorders. However, few studies have identified brain connectivity states that correlate with trait mindfulness in adolescence, nor have they assessed the reliability of such states.

**Methods:** To address this gap in knowledge, we rigorously assessed the reliability of brain states across 2 functional magnetic resonance imaging (fMRI) scan from 106 adolescents aged 12 to 15 (50% female). We performed both static and dynamic functional connectivity analyses and evaluated the test-retest reliability of how much time adolescents spent in each state. For the reliable states, we assessed associations with self-reported trait mindfulness.

**Results:** Higher trait mindfulness correlated with lower anxiety and depression symptoms. Static functional connectivity (ICCs from 0.31-0.53) was unrelated to trait mindfulness. Among the dynamic brains states we identified, most were unreliable within individuals across scans. However, one state, an hyperconnected state of elevated positive connectivity between networks, showed good reliability (ICC=0.65). We found that the amount of time that adolescents spent in this hyperconnected state positively correlated with trait mindfulness.

**Conclusions:** By applying dynamic functional connectivity analysis on over 100 resting-state fMRI scans, we identified a highly reliable brain state that correlated with trait mindfulness. The brain state may reflect a state of mindfulness, or awareness and arousal more generally, which may be more pronounced in those who are higher in trait mindfulness.

## Introduction

Adolescence is a time of rapid social, emotional, and brain maturation, and a critical time for the onset of mental illness. A meta-analysis of epidemiological studies found that 38% of adolescents with anxiety or fear-related disorders were diagnosed before the age of 14 (1). In the United States, 15% of adolescents experienced a major depressive episode in 2018 (2). Many adolescents continue to suffer from anxiety and/or depression into adulthood (3,4). There is a need to understand protective factors for mental illness in adolescence.

Trait mindfulness may be one such protective factor, defined as the tendency or disposition to pay attention to present moment experiences in a non-judgmental, accepting way (5). Trait mindfulness constructs were derived from mindfulness meditation practices and training (e.g., mindfulness-based stress reduction; (5)), which aim to not just cultivate mindfulness in the moment during meditation practice (‘state’ mindfulness) but extend the benefits to daily life (‘trait’ mindfulness). Trait mindfulness is typically measured using self-report scales which inquire about daily experiences. For example, the Child and Adolescent Mindfulness Measure (CAMM) has questions about emotional awareness: “I keep myself busy so I don’t notice my thoughts or feelings” (7). In adults, higher trait mindfulness scores have been consistently found to correlate with positive mental health outcomes, e.g. emotional well-being, and reduced psychopathology, e.g. reduced rumination and catastrophizing (8–10). In the last ten years, trait mindfulness scales have been validated for use with children and adolescents. These scales, e.g., the Mindful Attention Awareness Scale-Adolescents (11), and the Child and Adolescent Mindfulness Measure (CAMM) (7), have been found to likewise correlate with positive mental health status (12,13).

Despite the value of trait mindfulness for emotional well-being, little is known about the neural correlates of trait mindfulness in children and young adolescents (in contrast to numerous studies of adults; (14)).Three studies in children and adolescents have examined trait mindfulness correlates with task-based functional magnetic resonance imaging (fMRI activations) (15,16) and structural MRI (17). In consideration of traits that are supposedly consistent across contexts, resting-state fMRI is appealing because it measures functional connectivity of brain regions and networks that are independent of specific task demands. Resting-state fMRI contains both intrinsic, static components and time-varying, dynamic components (18–20). Static connectivity typically involves correlations between brain regions or networks over the course of an entire scan, whereas dynamic connectivity involves computing correlations within windows that are moved across the scan.

Only two studies have examined the relation of trait mindfulness to resting-state functional connectivity in children and adolescents. One study examined static connectivity in 23 adolescents who were remitted for MDD and 10 healthy controls, and found greater trait mindfulness to be inversely correlated with connectivity between the dorsolateral prefrontal cortex and the inferior frontal gyrus (two regions within the central executive network, CEN) (21). In the second study, we examined both static and dynamic resting-state functional network connectivity in relation to trait mindfulness in 42 children and adolescents (ages 6-17 years) with a focus on three brain networks: the CEN, the default mode network (DMN), and the salience network (SN) (22). Differing from the first study, we extracted functional networks using a group independent component analysis (ICA), which is a data-driven approach to finding spatially independent signals, each including a set of brain voxels sharing co-varying patterns (23). We found that more mindful children (as measured by the CAMM), showed *less* time in a brain state characterized by salience network (SN) anticorrelations with the other networks, a finding opposite from that found in adults (24). In addition, children who retrospectively reported more present-moment focused thoughts (less mind-wandering) during the scan showed less time in this brain state. Lastly, static functional network connectivity was not associated with trait mindfulness, suggesting the dynamic measures were more sensitive to trait mindfulness in this youth sample.

In the present study, we comprehensively investigated the functional brain bases of trait mindfulness in the largest sample (N>100) of adolescents to-date. We assessed the neural correlates of two trait mindfulness scales – the MAAS-A, which focuses on measuring day-to-day lapses of attention, and the CAMM, which focuses on emotional regulation and awareness. Our preregistered aim was to evaluate whether mindful adolescents show more or less time in an anticorrelated brain state, for instance, the DMN-SN anticorrelated brain state found in our previous study. We extracted functional brain networks using ICA, in keeping with evidence that ICA captures brain functional organization while retaining meaningful within-subject variability (25). We assessed 6 networks across the whole brain with both static and dynamic connectivity.

Prior to assessing correlations with mindfulness, we systematically investigated the scan-to-scan reliability of the functional connectivity measures. Reliability is an important component in brain-based individual differences research (26). If differences in functional imaging measures (e.g., connectivity) between individuals are not stable across imaging sessions, that is, lacking in consistency and/or agreement across sessions (which can be measured using intraclass coefficients (ICC) (27,28), then these measures cannot be predictively useful objective markers of traits of interest. A meta-analysis of static FC studies reporting reliability found ICCs for single connections were relatively low (29). Reliability issues could underlie discrepant results in studies of functional connectivity and trait mindfulness in adults (13). By focusing on reliability in this study, we hope to contribute to more replicable brain-behavior findings.

## Methods and Materials

### Preregistration

The analyses reported below were preregistered on the Open Science Framework (https://osf.io/wesu4).

### Participants

A total of 127 young adolescents (12.04-14.69 years) were recruited using social media, flyers, and local schools. Inclusion and exclusion criteria are found in **Supplement**. One hundred and six of the 127 participants (50% M) had at least one usable resting-state fMRI run (n=100 had 2 usable runs), the remainder were excluded for sleep (n=3), failure to complete the scan (n=12), or excessive head motion (n=6). Procedures were approved by the Massachusetts Institute of Technology Committee on the Use of Human Subjects. All adolescents and their legal guardians provided assent and consent. Parents and adolescents were compensated for their time.

**Table 1:** Adolescent demographics as reported by parents. Percentages are shown in parentheses. Income: Household Income (USD). M: million. ADHD: Attention Deficit Hyperactivity Disorder. NR: no response. A: ambidextrous.

|  |  |  |  |
| --- | --- | --- | --- |
| Variable | *N* = 106 | Variable | *N* = 106 |

| **Age (years)** |  | **Race/ Ethnicity (%)** |  |
| --- | --- | --- | --- |
| Mean | 13.46 | White | 65(61.3) |
| Range | 12.04-14.69 | Black | 8(7.5) |
| **Sex** |  | Asian | 4(3.8) |
| Male (%) | 53 (50%) | Hispanic | 10(9.4) |
| Female (%) | 53 (50%) | Pacific | 0(0) |
| **Handedness** |  | Amerindian | 0(0) |
| RH (%) | 92 (86.7%) | Other | 6(5.7) |
| LH (%) | 12 (11.3%) | Mixed | 13(12.3) |
| A (%) | 2 (1.8%) | **Pubertal Stage (%)** |  |
| **Income** |  | Pre | 1(0.9) |
| Mean (SD) | 137,742.20 \$ (138,169.90 \$) | Early | 8(7.5) |
| Range | 6500\$-1.25M \$ | Mid | 38(35.8) |
|  |  | Late | 16(15.1) |
|  |  | Post | 36(34.0) |
|  |  | NR | 5(4.7) |
|  |  | **Conditions (%)** |  |
|  |  | ADHD | 21(19.8) |
|  |  | Seizures | 2(1.9) |
|  |  | Concussions | 8(7.5) |
|  |  | Anxiety/Depression | 8(7.5) |
|  |  | On Medication | 25(23.6) |

### Self-report measures

Parents reported on household income, parental education, as well as their child’s race/ethnicity, medications, and medical conditions. Medications are listed in **Table S1**. Adolescents completed the remaining questions, including the self-report short pubertal scale (30), which provides two metrics: pubertal stage (categorical) and pubertal development (continuous).

We collected two trait mindfulness measures from the adolescents, the Mindful Attention and Awareness Scale for Adolescents (MAAS-A; hereafter ‘MAAS’) (11), and the Child and Adolescent Mindfulness Measure (CAMM) (7) (for details see **Supplement**). We collected the following additional self-report scales: depression (the MFQ; (31)), state and trait anxiety (STAI-C; (32)) and mind-wandering (the MWQ; (33)).

### Self-report analysis

We imputed missing item data (0.5% of responses) for the self-report questionnaires using predictive mean matching in the *mice* data package (34). There were no significant demographic differences between participants with usable and unusable rest data (**Supplement**). We also examined associations between participant demographics and mindfulness scores, to identify possible confounds before the brain imaging data analysis. Lastly, we examined the bivariate relationships between the self-report measures, to assess the external validity of the mindfulness questionnaires.

### Brain Imaging

#### Collection

Images were collected using a Siemens Prisma 3-T scanner with a 64-channel head coil. The protocol consisted of a T1-weighted scan, two fieldmaps (AP-PA and PA-AP), and then two resting-state scan runs (**Supplement**). Two resting-state scan runs of 4 minutes in duration each and TR of 0.8s were collected back-to-back. During the scans, participants were instructed to stare at a fixation cross. The T2*-weighted, gradient-recalled, multiband echo-planar imaging scan parameters were as follows for each of the two runs: multiband acceleration factor (8), TE (37 ms), flip angle (52°), echo spacing (0.58 ms), slice number (72), resolution (2 mm isotropic).

#### Preprocessing and denoising

Data were first preprocessed in fMRIPrep (v22.1.1), including T1 bias-field correction, fieldmap correction, brain extraction, normalization to the ICBM 152 nonlinear template, tissue segmentation, and motion correction procedures (35). We then spatially smoothed the functional data using a Gaussian kernel of 6 mm, and denoised the data in the CONN toolbox (36). We used standard denoising (37) (**Supplement)**, with the exception of despiking instead of removing high motion frames (18).

#### Network extraction and dynamic functional connectivity analysis

The procedure is outlined in **Figure 1**, and pipeline branches are shown in **Table 2**. We extracted networks using ICA, a data-driven analysis approach that finds spatially and temporally independent components (38). ICA was run on each individual separately, concatenating across the two runs, using FSL’s Melodic (39) (the *Individual ICA*), and on all participants together (the *Group ICA* in GIFT (v4.0c) (40) (**Supplement**). For Melodic, an automated network finding pipeline was executed with spatially cross-correlated networks from the 7-network Yeo atlas (41) and the components using FSL’s *fslcc* tool. As anticipated in our pre-registration, the limbic network often did not match any component closely (for ∼ 1 in 4 participants), whereas the other 6 networks were consistently found (See **Supplementary File 1** detailing the average correlations coefficients for each network). For this reason, the limbic network was omitted from analyses, resulting in 6 networks per participant: central executive network (CEN), dorsal attention network (DAN), default mode network (DMN), somatomotor cortex (SMC), ventral attention network (VAN), and visual network (VIS). Bilateral networks were treated as single networks (41). Network maps are provided in **Supplement,** as well as methods and network maps for Group ICA.

**Figure 1:**
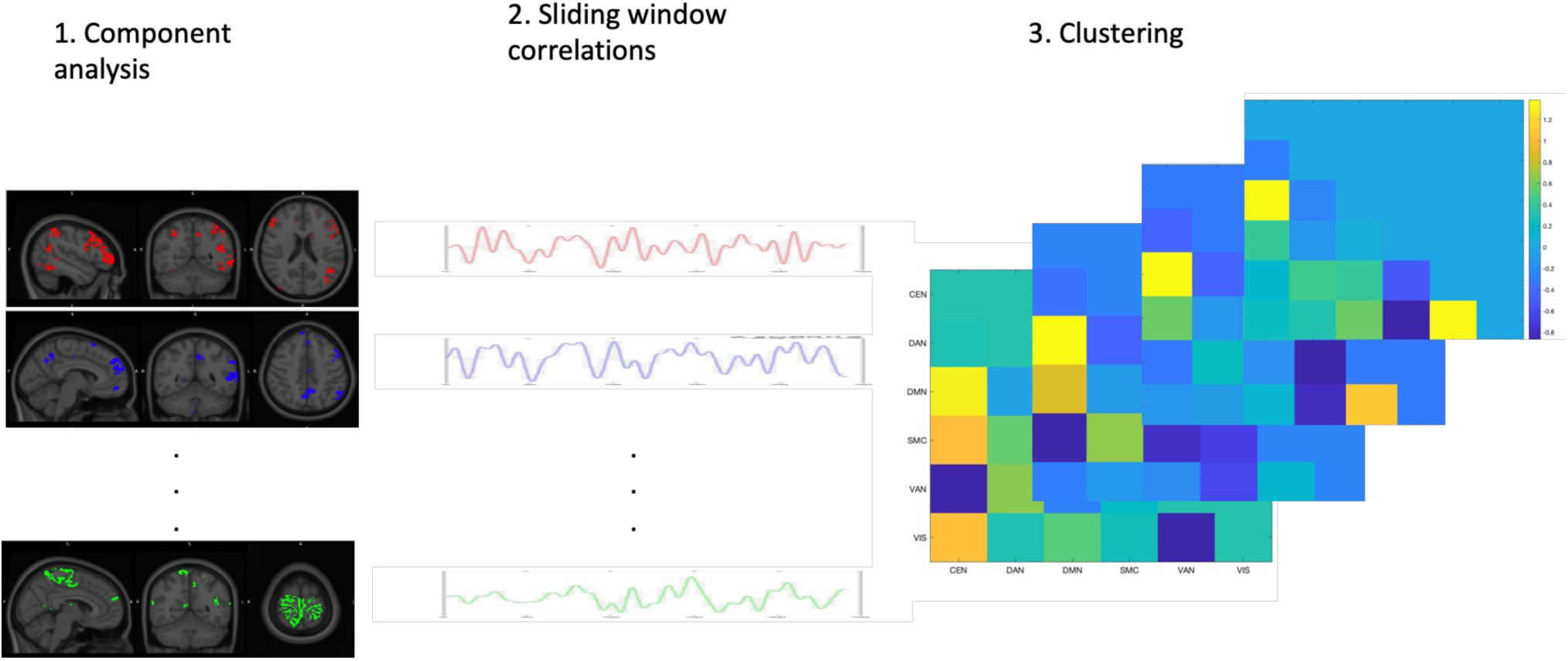
Dynamic functional connectivity analysis. In 1) functional networks are extracted from participants resting-state fMRI scans using ICA. In 2) timecourses of the functional networks are correlated within sliding windows to make connectivity matrices. In 3) the time-varying connectivity matrices are clustered into distinctive states using k-means.

We then extracted the mean time-courses of each network. First, we implemented a standard sliding window analysis using *icatb_compute_dfnc* from the GIFT toolbox (18). We took tapered windows with widths of 30 TRs (24 seconds) convolved with a Gaussian (3 TRs), and slid in steps of 1 TR (following our previous study (22)) resulting in 270 windows, to construct windowed correlation matrices (network connectivity, over time). We also explored a windowed approach without the Gaussian convolution. The windowed correlation matrices for all participants were then concatenated separately for run 1 and 2, resulting in two concatenated matrices (one for each run) that aggregated data across subjects.

To identify common patterns of connectivity (i.e., states) across participants and runs, k-means clustering was applied to these concatenated matrices in Matlab (scripts will be made available upon publication at https://osf.io/3gwt9/). We determined the optimal number of connectivity states by comparing the results from three separate cluster optimization techniques (**Supplement**).

#### Connectivity state analysis

Statistical analyses were performed using R. In rare cases, we removed connectivity states for signs of noise including extremely skewed distributions across participants (see Table 2), reclassifying windows using the remaining states. We calculated the proportion of time a participant spent in the connectivity state, the number of episodes in each state, and the average dwell time in each state, along with the total number of transitions. To examine the reliability of these dynamic measures, we conducted intra-class correlations (see below) across runs (e.g., does dwell time in state 1 in run 1 correlate with dwell time in state 1 in run 2?).

Finally, for those dynamic measures that were reliable, we calculated the average across the runs, and then assessed the relationships with the CAMM and MAAS scales individually. We controlled for framewise displacement, as well as pubertal stage (which we found correlated with mindfulness), given their possible impacts on functional connectivity (42–44). We controlled for multiple comparisons using false discovery rate (FDR) correction within each clustering solution for the set of reliable measures.

#### Static FNC analyses

Static functional network connectivity (sFNC) was calculated as the correlations between the networks across the entire time-courses (no windowing) of each run separately.

### Reliability

The two separate resting-state runs allowed assessment of the consistency of individual brain measures across runs. We implemented this using intra-class correlation coefficients (ICCs) for two dynamic measures: connectivity state dwell time and overall proportions of time, as well as for sFNC edges (network-network correlations). This reliability measure, specifically ICC(2,1) has been used previously in test-retest applications in brain imaging (45). In the case of comparisons of sFNC patterns across runs, we calculated edge-wise ICCs, and also a metric of pattern similarity (**Supplement**) (46).

## Results

### Self-Report and Demographics

We assessed relationships between demographics and the two mindfulness variables of interest, the MAAS (M = 68.46, SD = 13.03) and the CAMM (M= 31.25, SD = 4.66) questionnaires. The CAMM and the MAAS were positively correlated (*r* = 0.53), and both scales were negatively correlated with pubertal development (B = −1.91, *p* = 0.033 and B = −5.15, *p =* 0.049, respectively) when controlling for all other demographic variables. These relationships were not explained fully by age (**Tables S3 and S4**). As there was a relatively small range of ages (12–15) and a wider range of pubertal development, pubertal development was included as a covariate in further analyses.

In addition, the self-report measures showed a series of negative bivariate relationships between mindfulness and mental health symptoms and mind-wandering (see **Table S5)**.

### Static Functional Network Connectivity

We assessed correlations (sFNC) between the 6 networks selected in Individual ICA. ICC of the sFNC edges ranged from 0.31-0.53. When assessing the pattern similarity between connectivity matrices across runs, the average was *R* = 0.70. The z-scored static connectivity for Individual ICA is shown in **Figure 2A**. Static FNC was characterized by high VAN-SMC correlation, low positive DMN correlations with other networks, and moderately high DAN correlations with other networks. Group ICA resulted in similar reliabilities (**Supplement).** We assessed the relationships between the individual edges in the sFNC matrices (for both ICA methods) and trait mindfulness, and when controlling for multiple comparisons, none were significant (FDR-corrected *ps >* 0.25).

**Figure 2:**
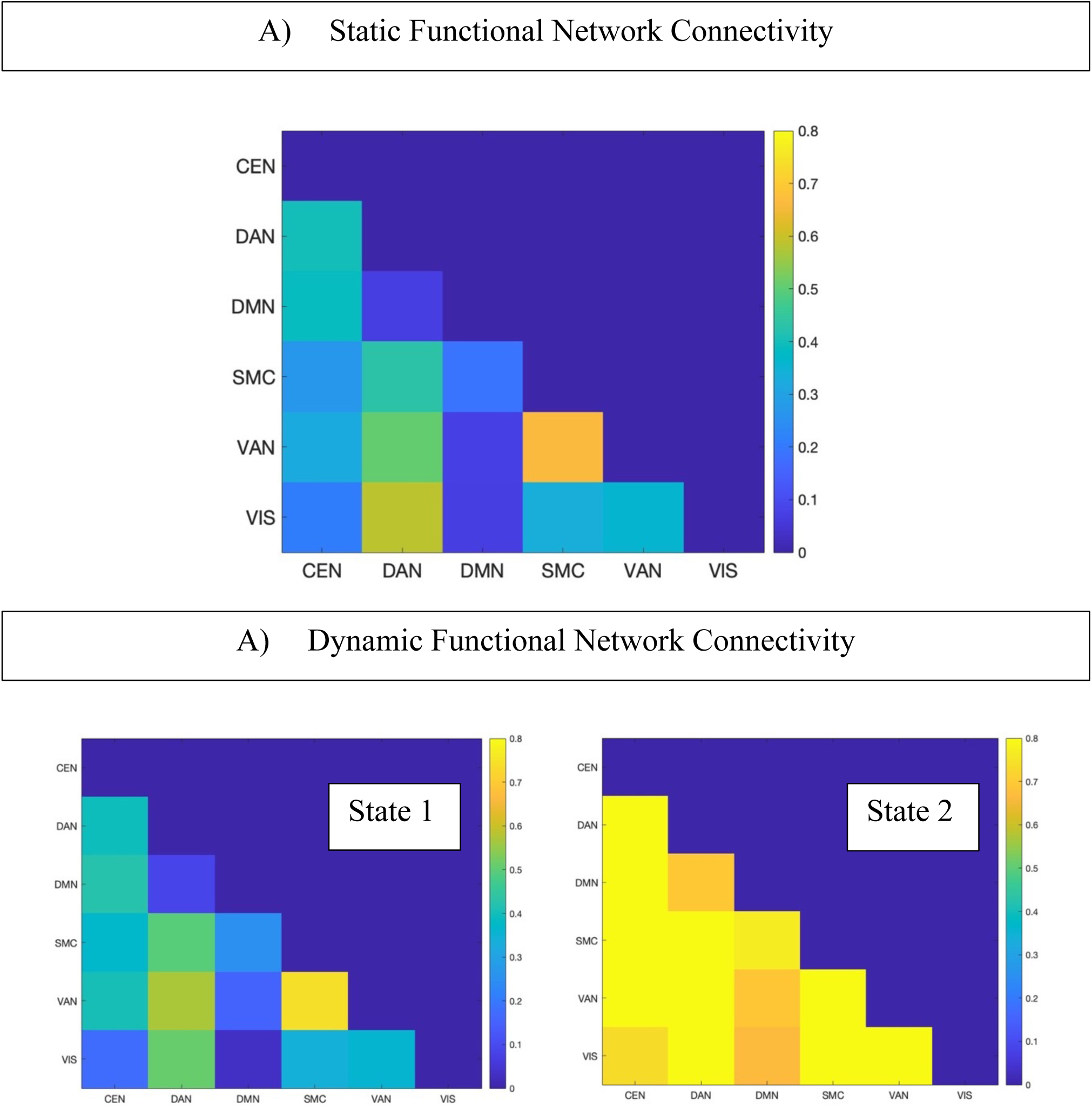
Static and dynamic functional connectivity matrices using the networks from individual ICA. Fisher’s z transform was applied to correlation coefficients. A) Static functional connectivity. B) Dynamic state 1 “static-like”, on left; Dynamic state 2: “hyperconnected”, on right. CEN: central executive network. DAN: Dorsal attention network. DMN: default mode network. SMC: sensorimotor cortex. VAN: ventral attention network. VIS: visual network.

### Dynamic Functional Connectivity

Reliability for each pipeline was estimated as a data-driven approach for finding candidates for the dynamic correlates of trait mindfulness. Reliabilities across runs for each combination of networks, convolution method, and optimal clusters are shown in **Table 2**. Generally, reliability fell in the poor to fair range, where 0–0.4 = poor, 0.4–0.6 = fair, 0.6–0.75 = good and 0.75–1 = excellent (27,28).

Reliabilities are shown for the dynamic ‘proportion of time’ metric, and not for dwell time, number of episodes, nor number of transitions, which almost always showed poor reliability. In general, reliability was higher for fewer states, for simple windowing without Gaussian convolution, and for individual ICA (which had fewer networks).

**Table 2:** Reliabilities of dynamic and static functional network connectivity. Details run1-run2 reliabilities of ‘proportion of time’ in brain states as well as sFNC strengths for different pipelines. NA: not applicable. Number of states vary based on clustering solutions (in parentheses are denoised selection). See methods for details on each branch. PS: pattern similarity, taking correlations between the sFNC patterns across runs.

| ICA | Sliding Window | Cluster Criterion | Number states | Reliabilities (ICC) |
| --- | --- | --- | --- | --- |
| Individual | No convolution | Cal. | 2 | 0.65 |
|  |  | Elbow | 4 | 0.23-0.38 |
|  |  | Gap | 9(7) | 0.27-0.54 |
|  | Convolution | Cal. | 2 | 0.37 |
|  |  | Elbow | 4(3) | 0.27-0.45 |
|  |  | Gap | 11(7) | 0.12-0.36 |
| Individual STATIC | NA | NA | NA | 0.31-0.53 (PS=0.70) |
| Group | No convolution | Cal. | 2 | 0.48 |
|  |  | Elbow | 4(3) | 0.24-0.39 |
|  |  | Gap | 11(5) | 0.07-0.33 |
|  | Convolution | Cal. | 2 | 0.33 |
|  |  | Elbow | 4 | 0.18-0.34 |
|  |  | Gap | 11(8) | 0.10-0.32 |
| Group STATIC | NA | NA | NA | 0.01-0.50 (PS=0.57) |

One solution fell into the ‘good’ range of reliabilities. That was individual ICA, using simple windowing, with two clusters. The clusters are shown in **Figure 2B**, and we term Brain State 1 the ‘static-like’ state (as it closely approximates the static functional connectivity), and Brain State 2 the ‘hyperconnected’ state with high correlations between all networks (Fisher’s *zs* > 0.6). Reliability of the proportion of time that the adolescents spent in each state was 0.65. Average dwell time in Brain State 1 showed a reliability of 0.56. Participants frequently switched between brain states, but tended to start in the hyperconnected state in both runs, and spent more time in the hyperconnected state overall (**Figure 3**).

**Figure 3:**
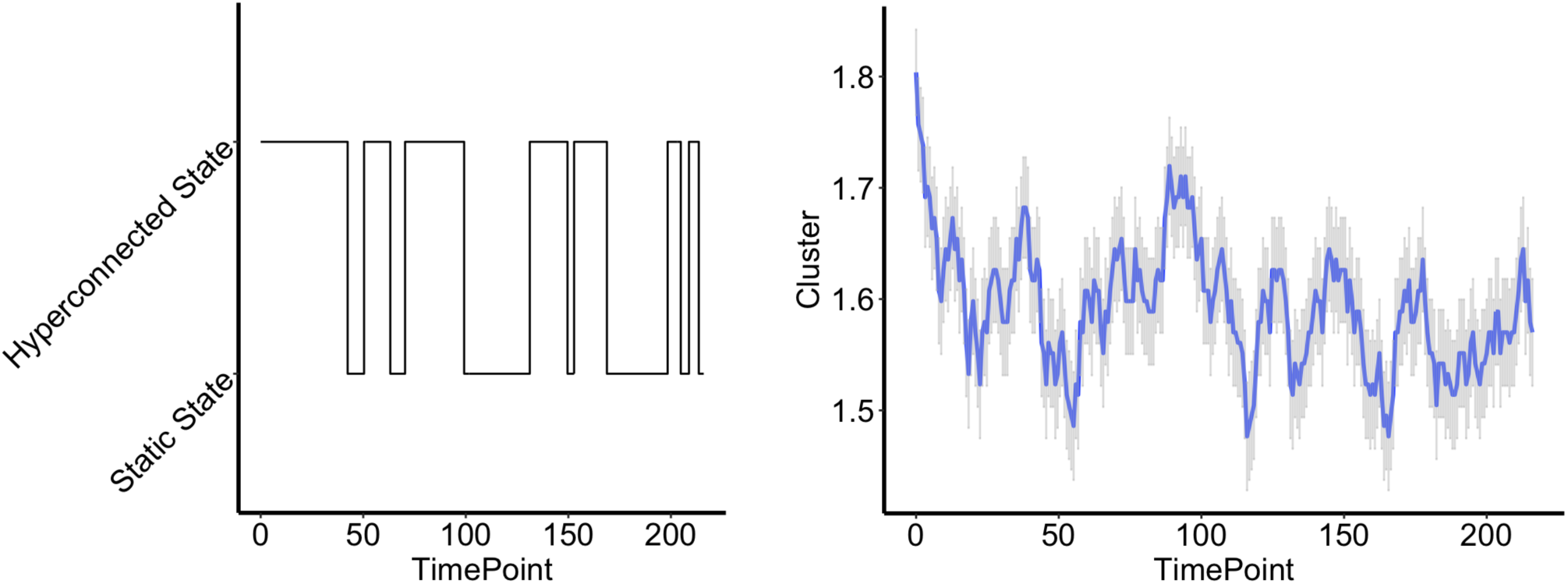
Timecourses of hyperconnected and static-like brain states. On left, a participant’s timecourse showing the switching between the static-like and hyperconnected brain states (1 and 2). On right, the timecourses are averaged across all participants for run 1. Higher values means more participants were in the hyperconnected brain state (2).

### Correlations with Trait Mindfulness

We then ran correlations with trait mindfulness for the reliable dynamic brain states, controlling for average framewise displacement and pubertal development. There was a significant positive relationship between the CAMM and the proportion of time spent in the hyperconnected brain state (B = 0.0076, *B* = 0.244, *p =* 0.018; **Figure 4**), which equates to a negative relationship with the proportion of time spent in the static-like brain state. When controlling for multiple comparisons between the two mindfulness questionnaires and the two reliable dynamic measures, the relationship was trend-level (*p* = 0.072). We ran the analyses without controlling for puberty, and found significant relationships for hyperconnected state and static-like state proportion time (B = 0.0079, *B* = 0.255, *p* = 0.008; B = −0.0079, *B =* −0.255, *p* = 0.008), and static-like dwell time (B = −0.29, *B* = −0.204, *p =* 0.036). When controlling for multiple comparisons, the relationship was significant for static-like and hyperconnected proportion time (*ps =* 0.033) and trend-level for static-like dwell time (*p* = 0.072). There were no significant relationships with the MAAS. In post-hoc exploratory analyses, we examined mental health symptoms, mind-wandering, and composites of the MAAS and CAMM and found no significant relationships to the brain states.

**Figure 4:**
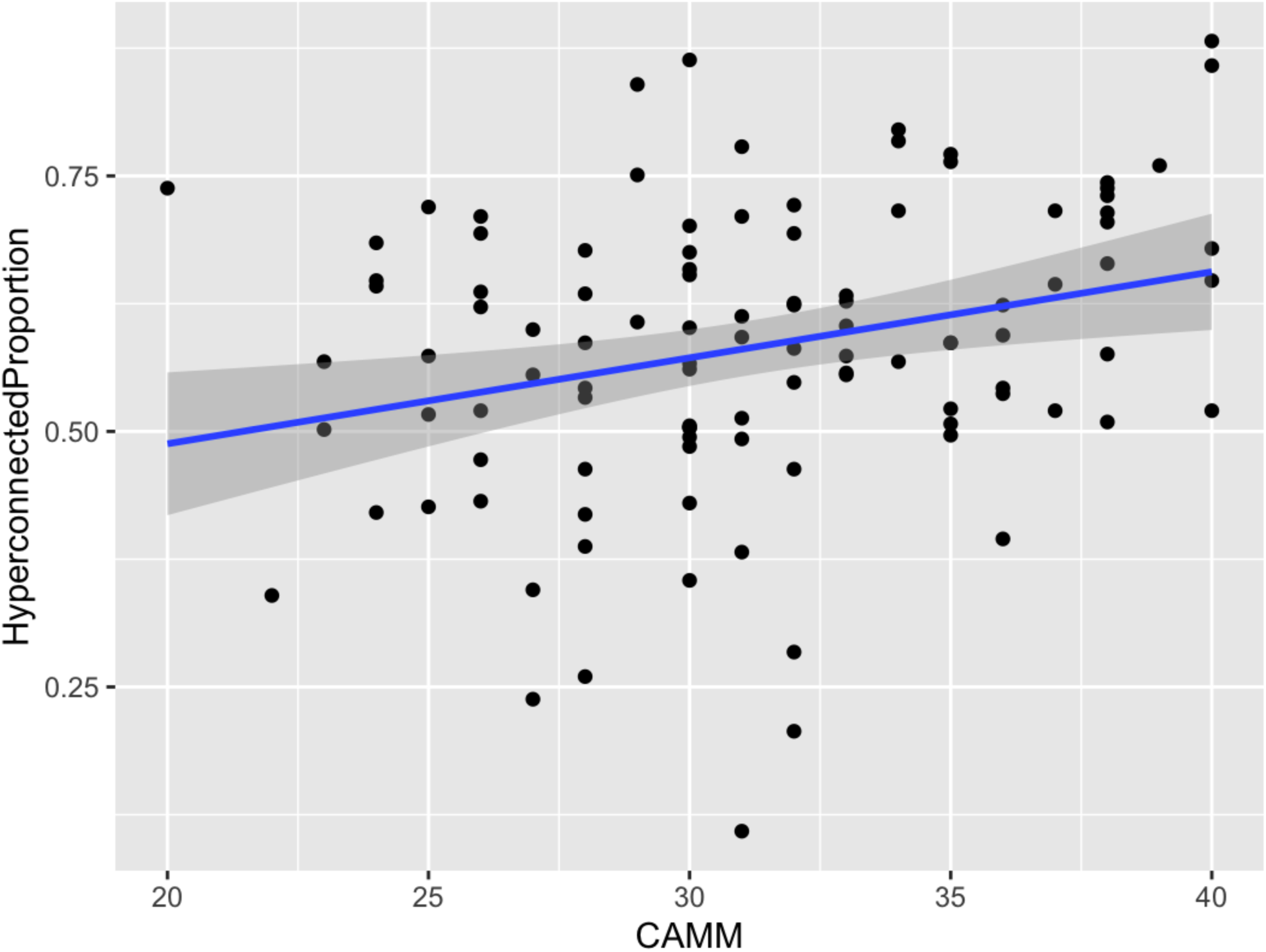
Brain dynamics correlate with trait mindfulness. CAMM: child and adolescent mindfulness measure. Hyperconnected Proportion (Brain State 2): Proportion of time in the hyperconnected brain state.

### Sensitivity Checks

There was no significant relationship between proportion of time in the two brain states and **average** framewise displacement (*p* = 0.15). We examined the relationship with instantaneous head motion. First, head motion was not higher at the beginning of runs as compared to the end (**Figure S3**). Second, there was no zero-order correlation between brain state and head motion. We ran a cross-correlation analysis, and found a significant relationship using bootstrapping between lags 10-21 across both runs, but coefficients were small (*R* < 0.05; **Figure S4)**. We additionally removed the first 5 timepoints of the runs, and the relationships between mindfulness and the brain states did not change. Robustness checks performed by removing high motion frames (scrubbing) did not change the pattern of results. When controlling for global static functional connectivity (averaged across all the edges), the relationship between hyperconnected proportion of time and mindfulness became trend-level (uncorrected *p* = 0.093).

### Other clustering solutions

As a post-hoc analysis, we examined whether there were relationships between mindfulness and the other clustering solutions with lower reliability. No relationships survived multiple comparisons.

Example connectivity states can be found in **Figures S5 and S6.** We examined connectivity states observed in our previous study and found no significant relationships with mindfulness (**Supplement).**

## Discussion

In this preregistered analysis, we investigated the resting-state functional magnetic resonance imaging (fMRI) correlates of trait mindfulness in 106 adolescents. We examined static functional connectivity between networks and applied dynamic functional connectivity methods to identify time-varying connectivity states, or ‘brain states’. Like our previous study with children and adolescents (22), we found no significant relationships between static functional connectivity and trait mindfulness. In terms of dynamic brain states, previous studies have suggested that brain states characterized by anticorrelations between attentional networks may be related to trait mindfulness, linked to more present-focused thinking and less mind-wandering (22,24,47). In our adolescent sample, we did not find any indication of this relationship. Instead, we found a positive correlation between trait mindfulness and time spent in a hyperconnected brain state: adolescents scoring higher on the CAMM measure spent more time in a state with elevated positive connectivity between all brain networks. This is the first observation of this relationship, and there are a number of reasons why this finding should be given consideration.

First, this study was, to our knowledge, the largest study relating trait mindfulness to functional connectivity in adolescents with over a hundred participants. We followed a preregistered analysis pipeline, which may help avoid concerns of false positives due to researcher degrees of freedom (48). In addition, we systematically explored different decision points in dynamic functional connectivity analysis. We examined networks extracted from ICA run separately on every individual compared to networks from Group ICA (23). We examined different windowing methods and different criteria for the clustering of connectivity matrices. We explored these decision points with the goal of optimizing reliability across runs of the resting-state data. Reliability is an important factor to consider in individual differences research (26). If the variance within-individuals is larger than the variance between-individuals, the measure may not be a good candidate to relate to a stable trait. Thus, we optimized reliability across resting-state runs before conducting correlations with mindfulness.

The two-cluster solution identified two reliable brain connectivity states – a ‘hyperconnected’ dynamic brain state, and a ‘static-like’ brain state, as it approximates the static functional connectivity over the course of the scans. The adolescents in this study tended to start in the hyperconnected brain state at the beginning of runs and spent more time in the state overall. Hyperconnected brain states have been reported previously (43,47,49–51), and so have dynamic brain states that are similar in network-network correlations to static functional connectivity (52). In addition, hyperconnectivity throughout many networks may be an emerging signature of the brain’s response to psychedelic drugs (53), and global hypoconnectivity has been found in depression (54,55). A common thread in these findings could be underlying cognitive flexibility and associated neural flexibility of brain states. Like psychedelics, mindfulness meditation may increase cognitive flexibility and lead to a wider range of brain states, including hyperconnected brain states (14,56). However, global hyperconnectivity has also been previously associated with head motion or physiological noise (57–59). Critically, we ruled out the possibility that head motion explained our findings by controlling for average head motion, and examining the relationship within individuals (where HM explained less than 0.25% of variance in brain states). Future studies should examine brain state relationships with physiological signals.

Previous neuroimaging studies have only examined single measures of mindfulness (21,22). We collected two self-report measures of mindfulness – the Child and Adolescent Mindfulness Measure (CAMM)(7), and the Mindful Attention and Awareness Scale – Adolescent (MAAS)(11). Both measures of mindfulness were negatively associated with depression, anxiety, and mind-wandering. The CAMM, but not the MAAS, was found to be correlated with the proportion of time participants spent in the hyperconnected brain state, and the average dwell time (how much time is spent on average per episode in the state). A meta-analysis of resting-state static functional connectivity and mindfulness interventions in adults found increased connectivity after mindfulness interventions, but it was specific to DMN-SN connections (60). The finding that mindfulness was positively related to the amount of time spent in greater connectivity across all examined (reliable) networks is a novel observation. This result is specific to the CAMM and no significant correlations were found when examining the MAAS, or composites of the MAAS and CAMM. Items on the CAMM were adapted from the Kentucky Inventory of Mindfulness Skills (KIMS; 61), specifically from three facets of the KIMS: *observing*, *acting with awareness*, and *accepting without judgment*. The CAMM has questions about emotional awareness, e.g. “I keep myself busy so I don’t notice my thoughts or feelings.” The CAMM shows negative correlations with rumination, stress, negative affect, and emotional and behavioral difficulties (7,12,62). The emotional focus of the CAMM contrasts with the ‘receptive attention’ focus of the MAAS (e.g. “I find myself doing things without paying attention”; 11). Thus our findings indicate that the hyperconnected brain state may be more frequent in individuals better able to notice and regulate their emotions, and contribute to emerging literature distinguishing different brain correlates of aspects of mindfulness (63,64,65).

### Limitations

As the hyperconnected state was more present in the beginning of scans, it is possible that it reflects an arousal response present when starting scans. Indeed, prior works have observed that moment-to-moment measures of physiological arousal fluctuate with greater global integration of functional connections (66). We caution attributing this finding to specific cognitive processes (19), as we did not collect any self-report measures about the scans from the adolescents. In addition, global static functional connectivity was correlated with time in the hyperconnected brain state, and the relationship to mindfulness was still present, but weaker, when controlling for global static FC. It is possible that the dynamic outcomes we derived are an index of hyperconnectivity more generally.

As described, it is unclear exactly what the hyperconnected brain state detected here and reported in prior studies reflects. In addition, one might question whether other, less reliable, brain states may be related to mindfulness. Researchers have cautioned against optimizing reliability at the cost of validity (67). Indeed, poor reliability does not eliminate the possibility of finding a relationship, but it puts an upper bound on the strength of the relationship (68). In a *post hoc* analysis, we examined whether any of the lower reliability brain states were related to mindfulness, and none were. It should also be noted that when we examined anticorrelated connectivity states previously identified in the literature (with high face validity), we found no relationship with mindfulness.

### Conclusions

We conducted the largest resting-state fMRI study of trait mindfulness in adolescents to-date, examining static and dynamic functional connectivity measures. We identified a reliable hyperconnected brain state that correlated with a trait mindfulness measure related to emotional responding. Future work could examine the state in a wider range of contexts including with momentary self-report assessments.

## Ethics statement

Procedures were approved by the Massachusetts Institute of Technology Committee on the Use of Human Subjects. All adolescents and their legal guardians provided assent and consent. Adolescents were compensated for their time.

## Data availability statement

Code and data will be uploaded to the following link upon publication: https://osf.io/3gwt9/.

