## Supplement for "Dynamic functional connectivity correlates of trait mindfulness in early adolescence"

**­­­­­­**

### Methods

#### Sample

This study was part of a larger project investigating factors influencing middle-school brain development and achievement. Primary inclusion criteria included seventh or eighth grade student at a local public school, English proficiency, and accompanying parent with English and/or Spanish proficiency. Primary exclusion criteria included MR contraindications, history of autism spectrum disorder or neurological disability, or premature birth (< 34 weeks).

#### Mindfulness questionnaires

The Mindful Attention and Awareness Scale for Adolescents (MAAS-A; hereafter ‘MAAS’) (1) consists of 15 items validated for an adolescent population, with a focus on attentional lapses, e.g., “I find it difficult to stay focused on what's happening in the present.” In this sample, the Cronbach’s Alpha of the MAAS was 0.91. The other questionnaire was the Child and Adolescent Mindfulness Measure (CAMM) (2). The CAMM consists of ten items validated for children and adolescents, with a broader focus than the MAAS, e.g.,“I tell myself that I shouldn't feel the way I'm feeling.” In this sample, the Cronbach’s Alpha of the CAMM was 0.81. We examined both scales individually given their theoretical distinctions (3).

#### Usable Resting-state Analyses

We examined whether there were any differences between those participants with usable resting-state data and non-usable resting-state data using logistic regressions. There were no systematic differences in demographics between adolescents with usable and without usable rest data (*ps* > 0.35; See **Table S2**). However, adolescents with usable resting-state data had lower state anxiety than those without usable data (M = 30.4 vs M = 32.9, respectively, *p* = 0.040).

#### Structural brain imaging:

One high-resolution, multiecho, magnetization-prepared rapid gradient echo T1w image was acquired along with an additional vNav setter for prospective motion correction. The vNav-enabled scan estimated motion throughout the T1w scan and reacquired/replaced k-space data unduly affected by motion (4). T1w scans featured a 0.8-mm isotropic voxel size with 320 slices, acquired in the sagittal orientation, repetition time (TR)/echo time (TE) = 4000/1.06 msec.

#### Denoising:

In particular, we applied linear regression to remove the following parameters from each voxel: (a) 5 noise components each from minimally-eroded WM and CSF (one-voxel binary erosion of voxels with values above 50% in posterior probability maps), respectively, based on aCompCor procedures (5) (b) 12 motion parameters (3 translation, 3 rotation, and associated first-order derivatives); (c) linear BOLD signal trend within session. We did not apply global signal regression. In a separate step after nuisance regression, data was temporally filtered with a bandpass of 0.01–0.1 Hz. Rather than removing high motion frames, which may be non-optimal for dynamic functional connectivity (6), we conducted despiking after regression. Despiking involved a tangent squashing function, which was linear for signals within ± 3SD of the mean, but applied an exponential function outside to minimize undue effects of outliers.

#### ICA:

##### *Overview*

ICA was run on each individual separately using FSL’s Melodic (7) (the *Individual ICA*), and on all participants together (the *Group ICA* in GIFT (v4.0c) (8). Different software packages were used because the network classification scripts were developed separately (specifically, the matching of Melodic components to networks). We initially estimated networks separately by run as noted in the preregistration, but found low reliability of the proportion of time in dynamic brain states across runs, perhaps because more data are required for robust estimates (see (9–11) demonstrating better functional connectivity reliability with longer scans).

##### *Individual ICA*

For Individual ICA, we ran Melodic on each participant separately, by first concatenating the two resting-state runs with dimensionality estimation using the Laplace approximation to the Bayesian evidence of the model. This resulted in a mean of 19.33 components per individual (range 15 to 24). Then, an automated network finding pipeline was executed with spatially cross-correlated networks from the 7-network Yeo atlas (12) and the components using FSL’s *fslcc* tool*,* identifying a single most correlated match for each of the networks. As anticipated in our pre-registration, the limbic network often did not match any component closely (for ~ 1 in 4 participants), whereas the other 6 networks were consistently found (See **Supplementary File 1** detailing the average correlations coefficients for each network). For this reason, the limbic network was omitted from analyses, resulting in 6 networks per participant: central executive network (CEN), dorsal attention network (DAN), default mode network (DMN), somatomotor cortex (SMC), ventral attention network (VAN), and visual network (VIS). Bilateral networks were treated as single networks (Yeo et al., 2011). Network maps are provided in **Figure S8**.

##### *Group ICA*

For Group ICA analysis, we first conducted two PCA data reduction steps using SVD estimation, and then derived 12 components using the Infomax algorithm, repeated 20 times in ICASSO to estimate stable components (13). We conducted semi-automated network selection using the Yeo atlas as a reference. We also included the salience network (SN) from the GIFT toolbox atlas (*RSN)* in our analysis, which is not included in the Yeo atlas. This was justified given previous literature focusing on the SN, as well as the ability when doing semi-automated classification to distinguish the SN from the VAN. We took the 12 components identified by the ICA and conducted spatial cross-correlations. Two authors then independently labelled the components while blinded to the components’ cross-correlations. Finally, the networks were selected by the lead author based on the best correspondence between authors and the calculated cross-correlations. We applied this semi-automated method because the number of components to assess was small compared to the >2000 components in the individual ICA. It was found that some of the networks were fractionated, for example left and right CEN, in which case both components were included. We again left out the limbic network, resulting in 10 networks, rCEN, lCEN, DAN, pDMN, aDMN ,SAL,SMC, VAN, VIS1, VIS2). Network maps are provided in **Figure S9**.

#### Clustering connectivity states

The algorithms consisted of (1) the Calinski-Harabacz algorithm (implemented in Matlab) (2) the elbow criterion (implemented using the *icatb_optimal_clusters* function from the GIFT toolbox) (3) the gap criterion (also implemented using *icatb_optimal_clusters)*. These methods evaluate the quality of the clustering by comparing distances within-clusters to distances between-clusters, with the goal of having low within-cluster distances and high between-cluster distances. After determining the optimal number of clusters using these methods (generally, 2 for Calinski, 4 for elbow, >6 for Gap), we reran k-means using the *icatb_kmeans_clustering* function from the GIFT toolbox. The clusters were compared across runs using correlations to compare the patterns over the network-network edges and the lead author’s judgement. The clusters are hereafter called connectivity states.

#### Pattern similarity of connectivity states

Pattern similarity was evaluated by taking within-individual correlations between the sFNC patterns across runs, averaging across participants (using fisher’s *z*), and then converting back to an average *R.*

#### Exploratory analyses

We conducted non-preregistered analyses to better explain and interpret the dynamic functional connectivity results. We examined the relationship between instantaneous head motion and the brain connectivity states in the two-state solution using cross-correlations. Further, we examined sFNC between the networks. We investigated the reliability of the sFNC estimates (as a reliability benchmark), assessed whether they were correlated with mindfulness (to assess the sensitivity of dynamic vs static), and included global sFNC as a control variable in the regressions between the dynamic measures and mindfulness.

### Results

#### Puberty

The magnitude of the correlation without controlling for other variables was *r* = -0.34 for the CAMM, and *r =* -0.27 for the MAAS (see **Figure S1**).

#### Mindfulness correlations with other measures

The CAMM and MAAS scales were significantly negatively correlated with depression symptoms (*r* = -0.37, *r* = -0.53), mind-wandering (*r* = -0.41, *r* = -0.63), perceived stress (*r* = -0.40, *r* = -0.50), as well as trait (*r* = -0.44, *r* = -0.60) and state anxiety (*r* = -0.31, *r* = -0.34).

#### sFNC for group ICA

Reliabilities were similar when examining the static FNC using the group networks from ICA; ICC of individual edges ranged from 0.01-0.50, and average pattern similarity was *R* = 0.56. Static FNC for Individual ICA was uniformly positive, whereas there were anticorrelations in Group ICA (**Figure S2**); however, these anticorrelations were isolated to connections involving the anterior DMN, which was averaged with posterior DMN in Individual ICA.

#### Attempted replication

We also examined the four group networks used in our previous study (13), constraining the number of clusters to 5 as per the previous analysis. We did see some correspondence between clusters across the two datasets and sites (3/5 clusters; **Figure S7)**. For example, there was a SN anticorrelated state. Reliability was poor-to-fair for the proportion-of-time measures, but near-zero for number of transitions. We failed to observe a relationship between mindfulness and the proportion of time spent in the SN anticorrelated state (CAMM *p* = 0.977, MAAS *p* = 0.717), nor in other states (uncorrected *ps* > 0.06)*.*

### Tables and Figures

| Medication type | Quantity of participants |
| --- | --- |
| ADHD | 16 |
| Anxiety | 6 |
| Asthma or Allergy | 3 |
| Mood | 5 |
| Other | 6 |

**Supplemental Table 1:**

Medications taken by adolescents in study. Some participants took more than one.

|  | Estimate | Standard Error | z value | Pr(>\|z\|) |
| --- | --- | --- | --- | --- |
| (Intercept) | 2.583 | 10.362 | 0.249 | 0.8032 |
| WhiteOrNot | -0.646 | 1.241 | -0.520 | 0.6030 |
| Age | -0.095 | 0.848 | -0.113 | 0.9103 |
| `Pubertal Development Scale` | 0.411 | 1.080 | 0.380 | 0.7037 |
| Sex | 1.009 | 1.244 | 0.811 | 0.4171 |
| ZAvg_SES | -0.110 | 0.550 | -0.200 | 0.8414 |
| **Supplemental Table 2:** Results of logistic regression predicting usable rest data based on demographics. WhiteOrNot: binary variable corresponding to race of white or not. ZAvg_SES: a composite of maternal education and household income. \| Estimate \| Standard Error \| t value \| Pr(>\|t\|) \|; (Intercept) \| 54.387 \| 8.503 \| 6.396 \| 0.0000 \| ***; Pubertal.Development.Scale \| -1.806 \| 0.774 \| -2.335 \| 0.0214 \| *; Age \| -1.354 \| 0.699 \| -1.938 \| 0.0553 \| .; *Signif. codes: 0 <= '***' < 0.001 < '**' < 0.01 < '*' < 0.05* \| \| \| \| \| **Supplemental Table 3:** Results of linear regression model predicting child and adolescent mindfulness measure (CAMM) by pubertal development and age. | | | | |

|  | Estimate | Standard Error | t value | Pr(>\|t\|) |  |
| --- | --- | --- | --- | --- | --- |
| (Intercept) | 110.422 | 24.674 | 4.475 | 0.0000 | *** |
| Pubertal.Development.Scale | -4.280 | 2.246 | -1.906 | 0.0594 | . |
| Age | -2.245 | 2.028 | -1.107 | 0.2706 |  |
| *Signif. codes: 0 <= '***' < 0.001 < '**' < 0.01 < '*' < 0.05* | | | | | |

**Supplemental Table 4:**

Results of linear regression model predicting MAAS by pubertal development and age.

| Measure | CAMM | MAAS | MFQ | MWQ | PSS | Anxiety state |
| --- | --- | --- | --- | --- | --- | --- |
| CAMM |  |  |  |  |  |  |
| MAAS | 0.53**** |  |  |  |  |  |
| MFQ | -0.37**** | -0.53**** |  |  |  |  |
| MWQ | -0.41**** | -0.63**** | 0.40**** |  |  |  |
| PSS | -0.40**** | -0.50**** | 0.48**** | 0.42**** |  |  |
| Anxiety state | -0.31** | -0.34*** | 0.45**** | 0.25* | 0.49**** |  |
| Anxiety trait | -0.44**** | -0.60**** | 0.66**** | 0.49**** | 0.63**** | 0.50**** |

**Supplemental Table 5:**

Relationships between self-report scales. CAMM: Child and Adolescent Mindfulness Measure. MAAS: Mindful Attention and Awareness Scale- Adolescents. MFQ: Moods and Feelings Questionnaire. Likely depressed: MFQ scores > 27. MWQ: Mind-wandering Questionnaire. PSS: Perceived Stress Scale. Anxiety state: STAI-C subscale. Anxiety trait: STAI-C subscale. *: *p* < 0.05, ** *p* < 0.01, *** *p* < 0.001, **** *p* < 0.0001

NR

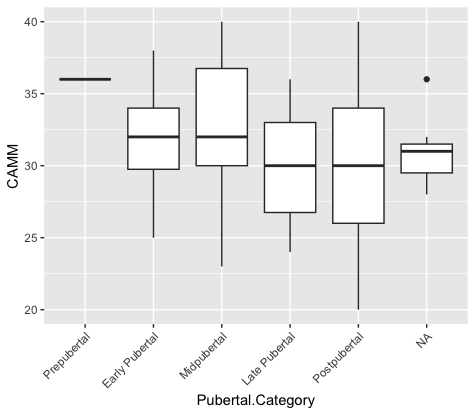

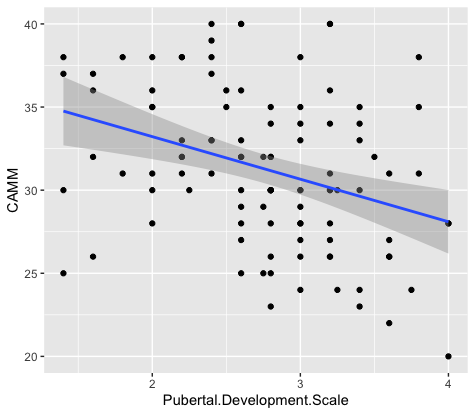

**Supplemental Figure 1:**

Relationships between the Child and Adolescent Mindfulness Measure (CAMM) and short self-report pubertal development scale. In A) the pubertal scores provide categorical cutoffs, in B) they are summed for a continuous measure. NR: no response.

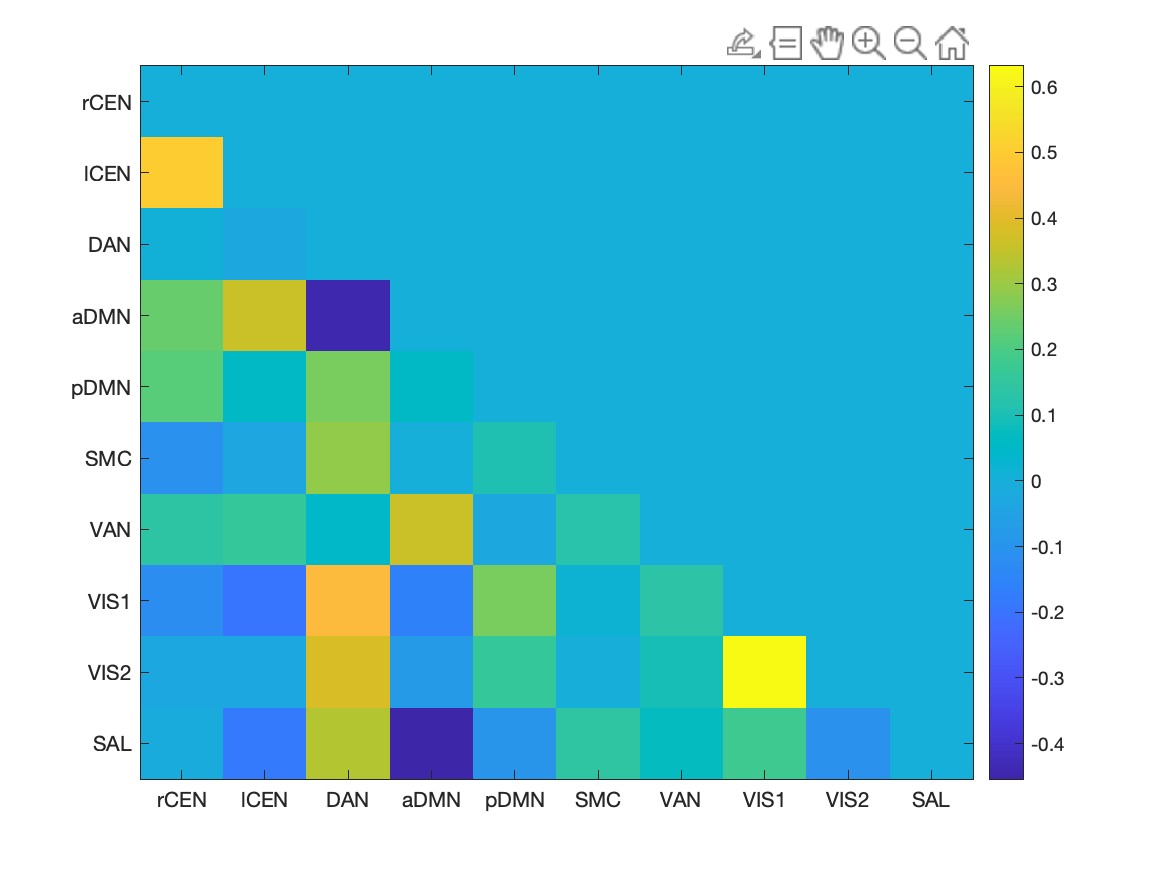

**Supplemental Figure 2:**

Static functional connectivity for the 10 networks identified by GroupICA. rCEN: right central executive network. lCEN: left central executive network. DAN: dorsal attention network. pDMN: posterior default mode network. aDMN: anterior default mode network. SMC: sensorimotor cortex. VAN: ventral attention network. VIS1: visual network 1 (primary). VIS2: visual network 2 (extrastriate). SAL: salience network.

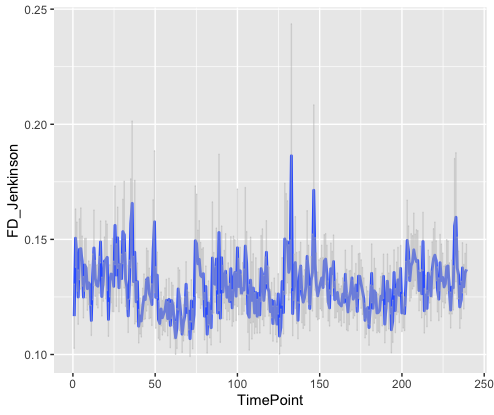

**Supplemental Figure 3:**

Instantaneous head motion (using the Jenkinson method) averaged across participants over the course of run 1.

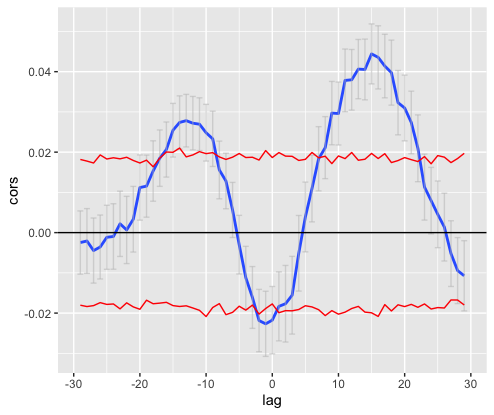

**Supplemental Figure 4:**

Cross-correlations between brain state and instantaneous head motion. Lags are shown on the x-axis, and correlation coefficients on the y-axis. Blue lines represent average correlation over participants, with grey standard error bars. Red lines represent bootstrapped intervals calculated by doing 1000 permutations of the brain state data.

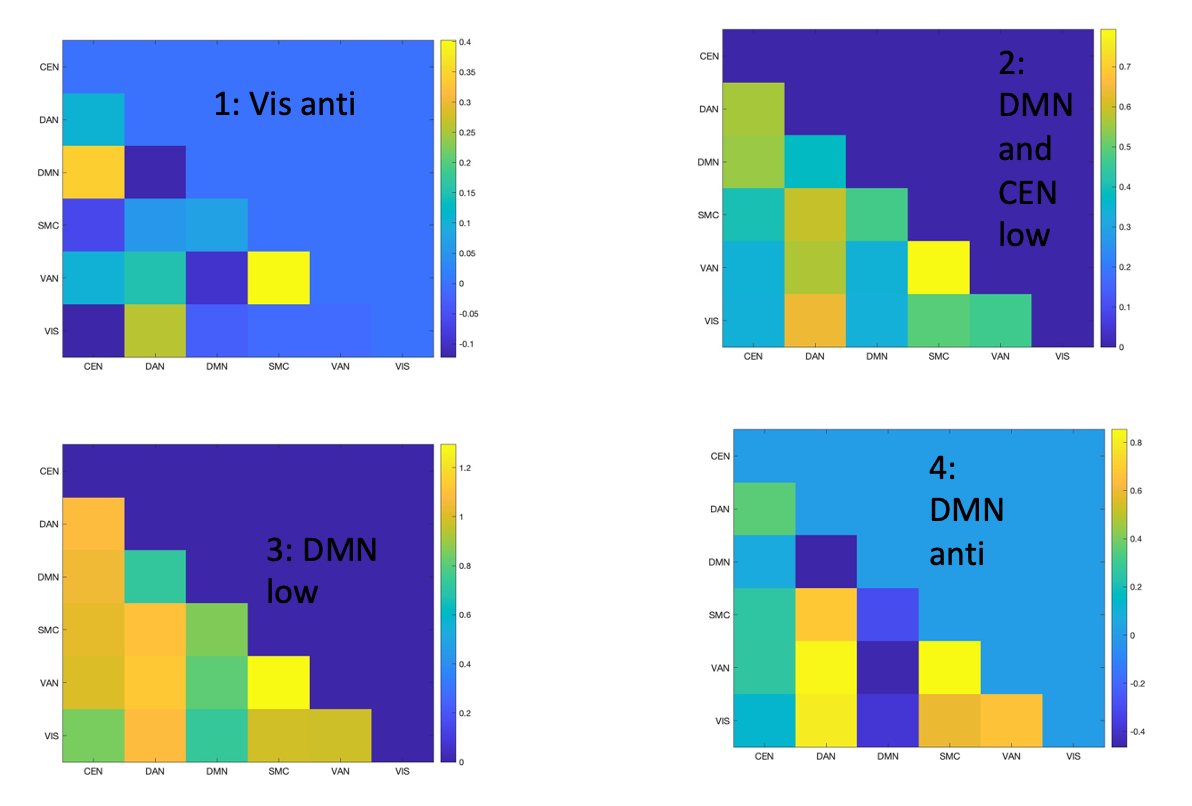

**Supplemental Figure 5:**

Dynamic brain states found using individual ICA, convolution, and the elbow method. CEN: central executive network. DAN: Dorsal attention network. DMN: default mode network. SMC: sensorimotor cortex. VAN: ventral attention network. VIS: visual network.

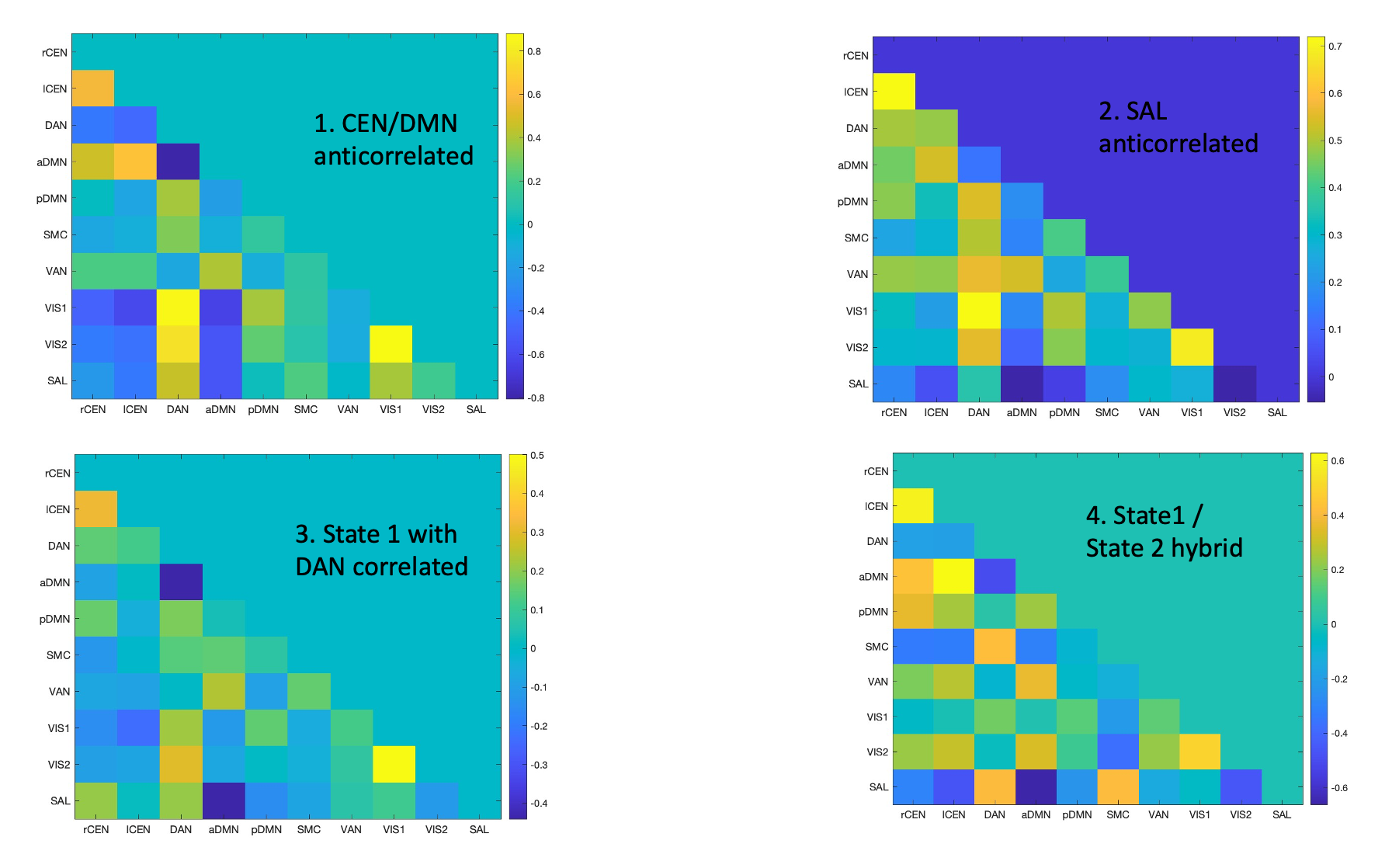

**Supplemental Figure 6:**

Dynamic brain states found using group ICA, convolution, and the elbow method. rCEN: right central executive network. lCEN: left central executive network. DAN: dorsal attention network. pDMN: posterior default mode network. aDMN: anterior default mode network. SMC: sensorimotor cortex. VAN: ventral attention network. VIS1: visual network 1 (primary). VIS2: visual network 2 (extrastriate). SAL: salience network.

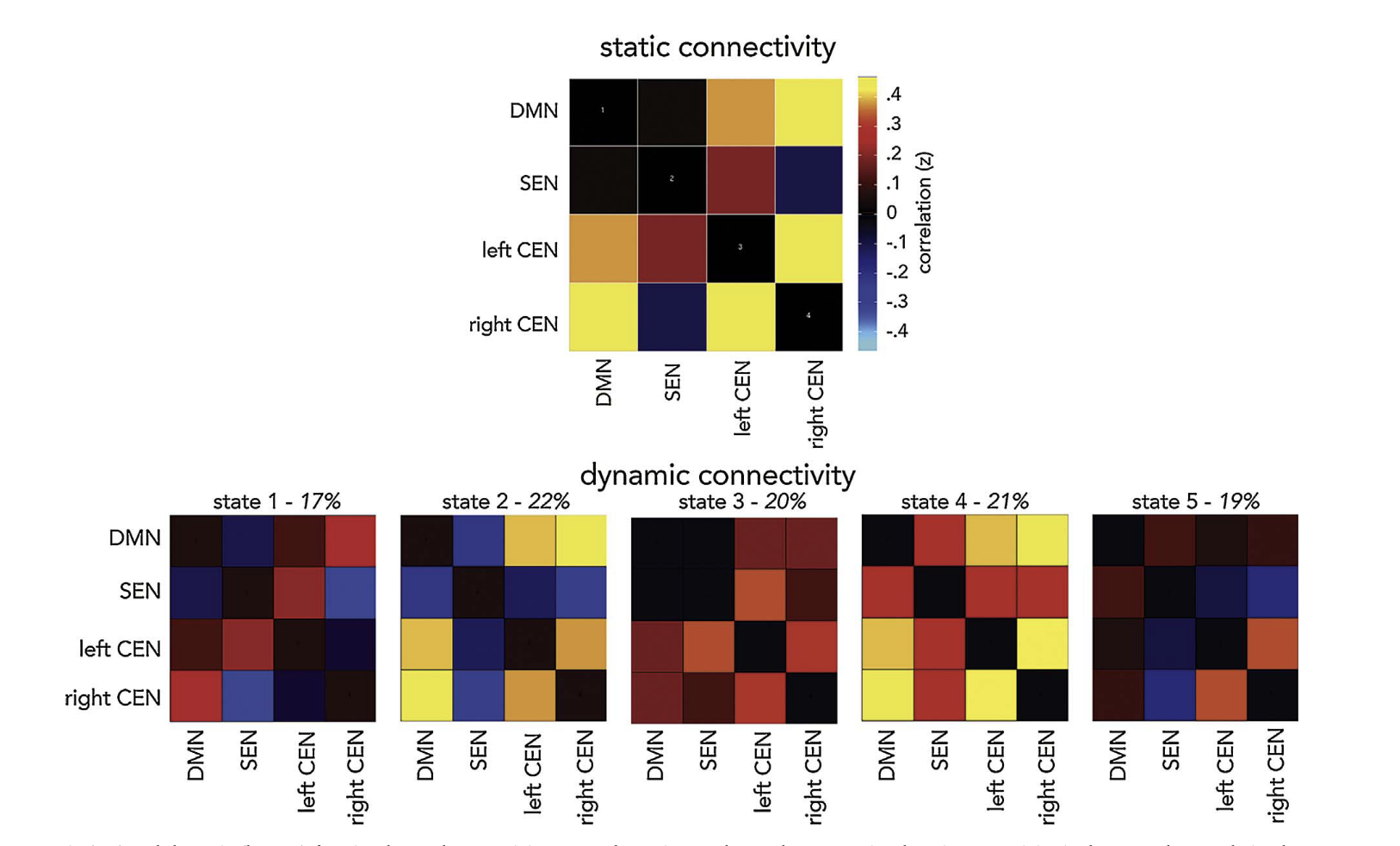

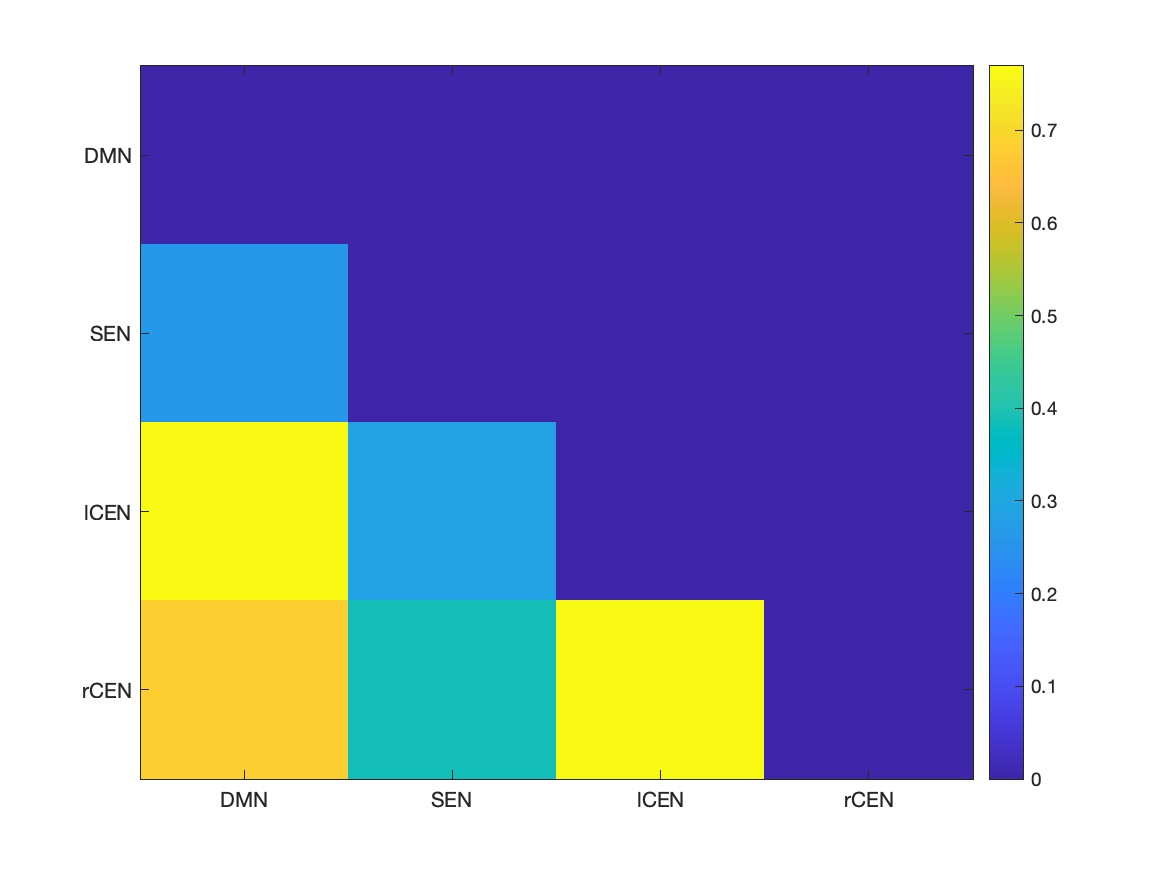

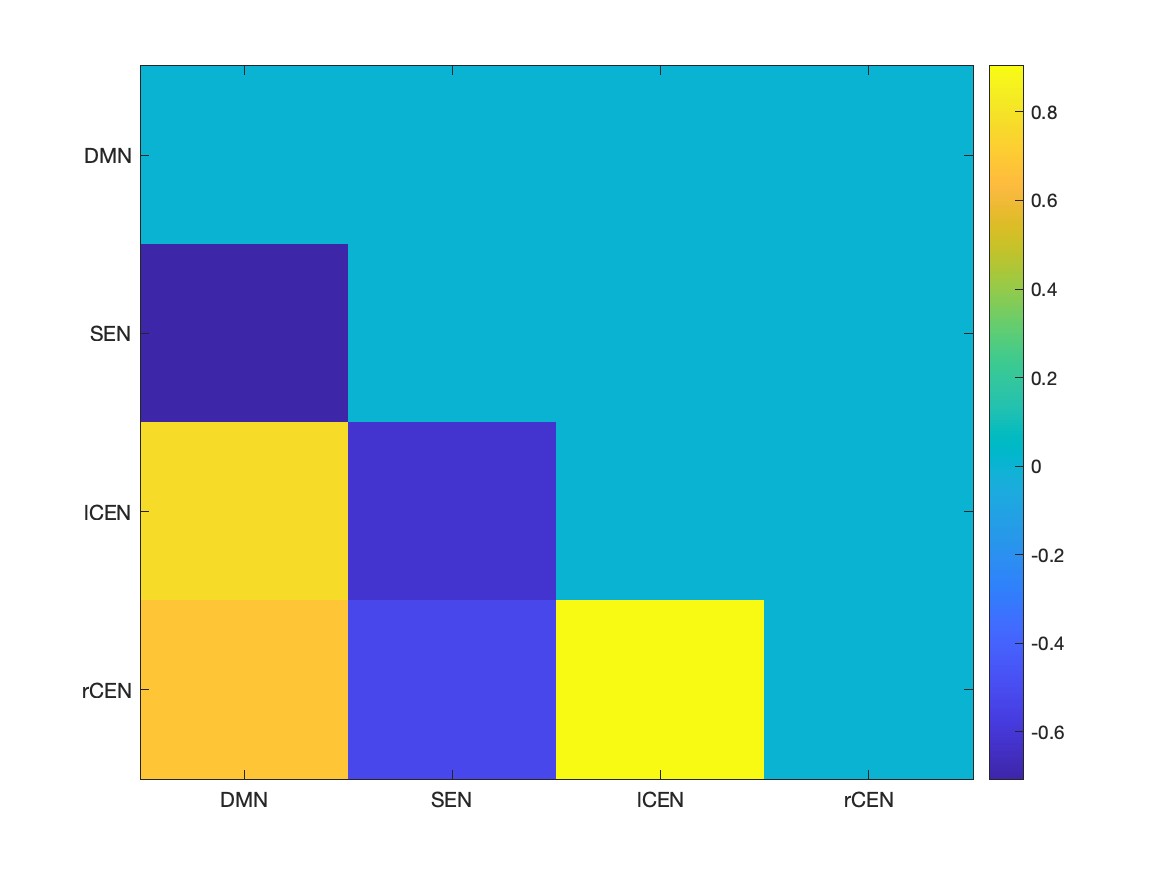

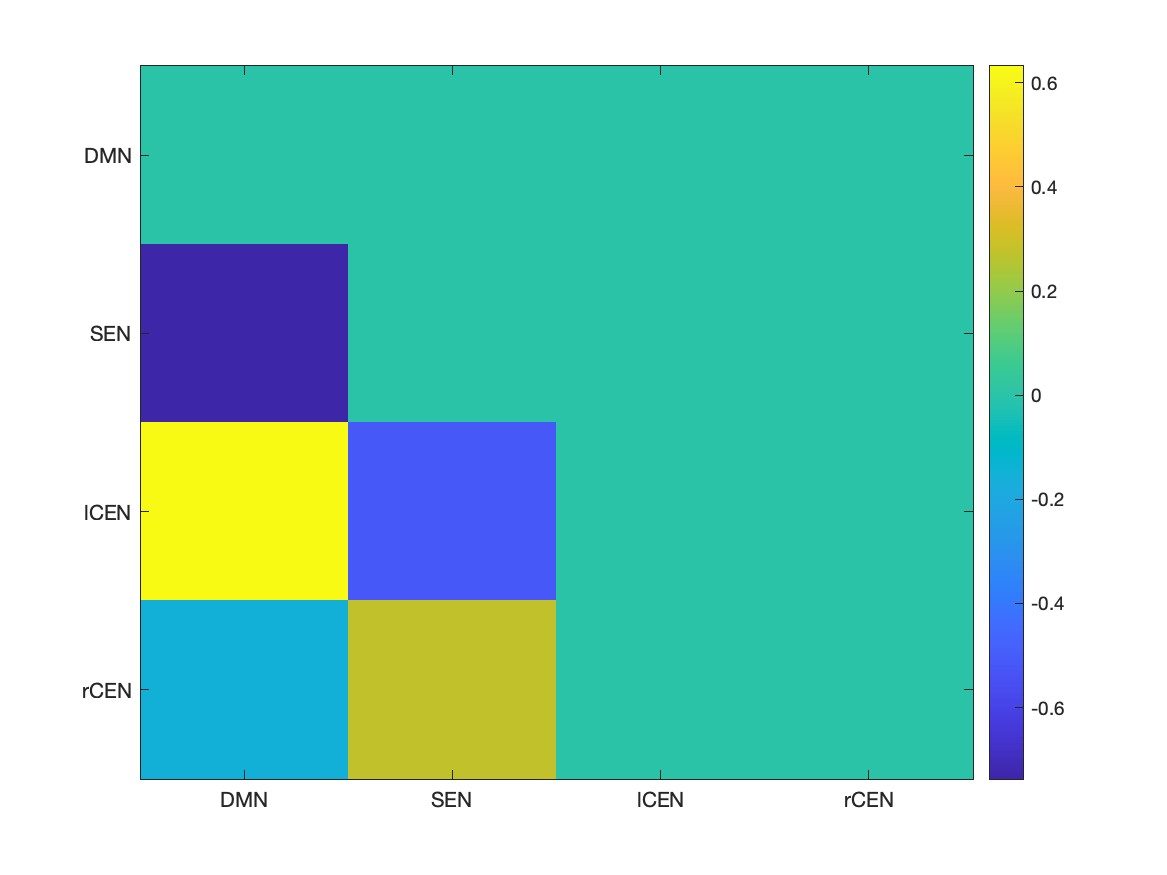

**Supplemental Figure 7:**

Comparison of dynamic brain states between Marusak et al. 2018 and the present study. On top, the five states observed by Marusak et al., 2018. SEN: Salience Network, CEN: central executive network (analogous to FPN). On bottom, three of the five states we observed that match those states. Specifically, we observed a SEN/ CEN correlated brain state, a SEN anticorrelated brain state, and a low SEN brain state.

**Supplementary Figure S8:** Brain networks from individual ICA

**CEN**

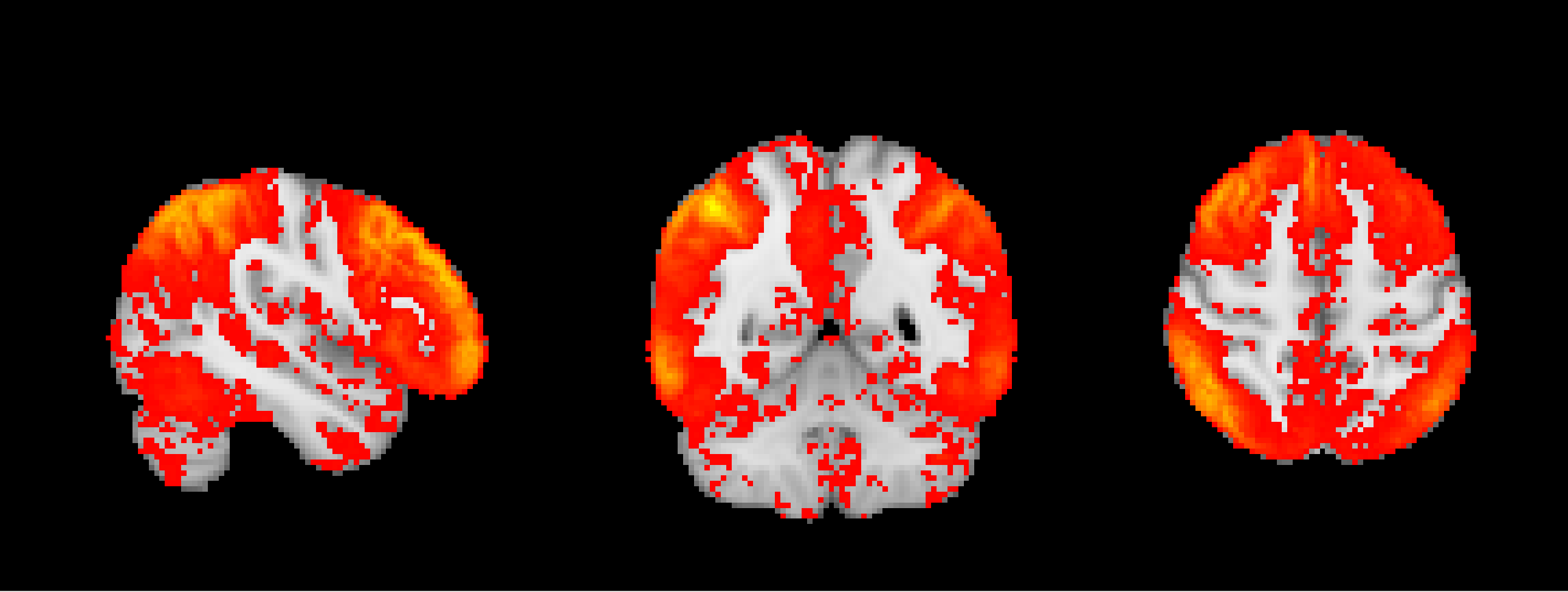

Z= 56 mm

X= 48 mm

Y= -52 mm

**DAN**

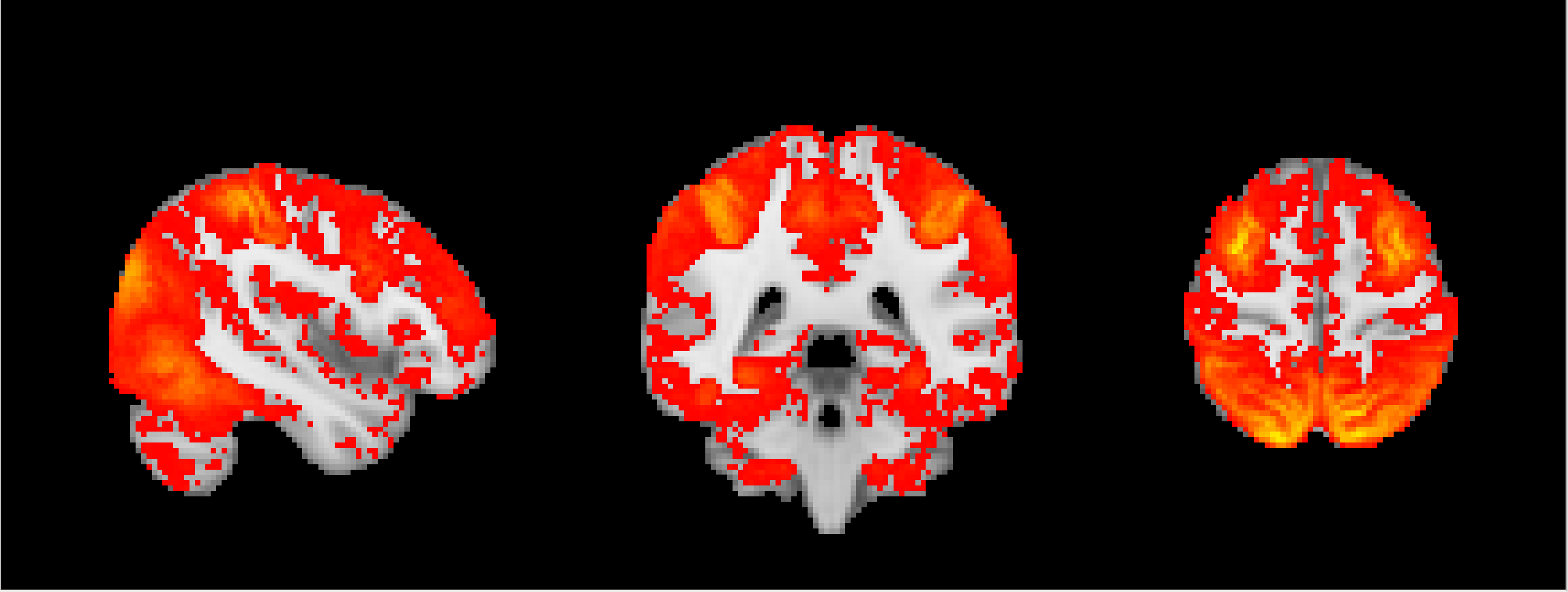

Z= 62 mm

Y= -40 mm

X= 45 mm

**DMN**

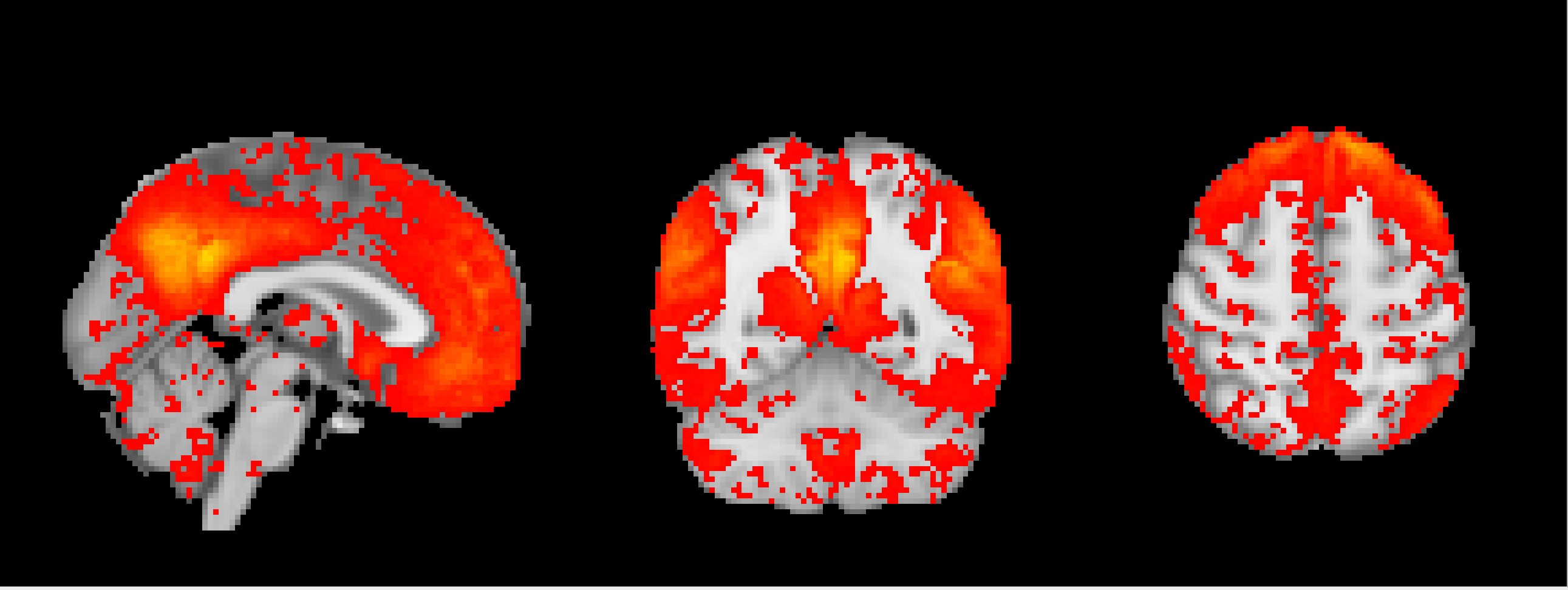

X= 0.5 mm

Y= -56 mm

Z= 56 mm

**SMC**

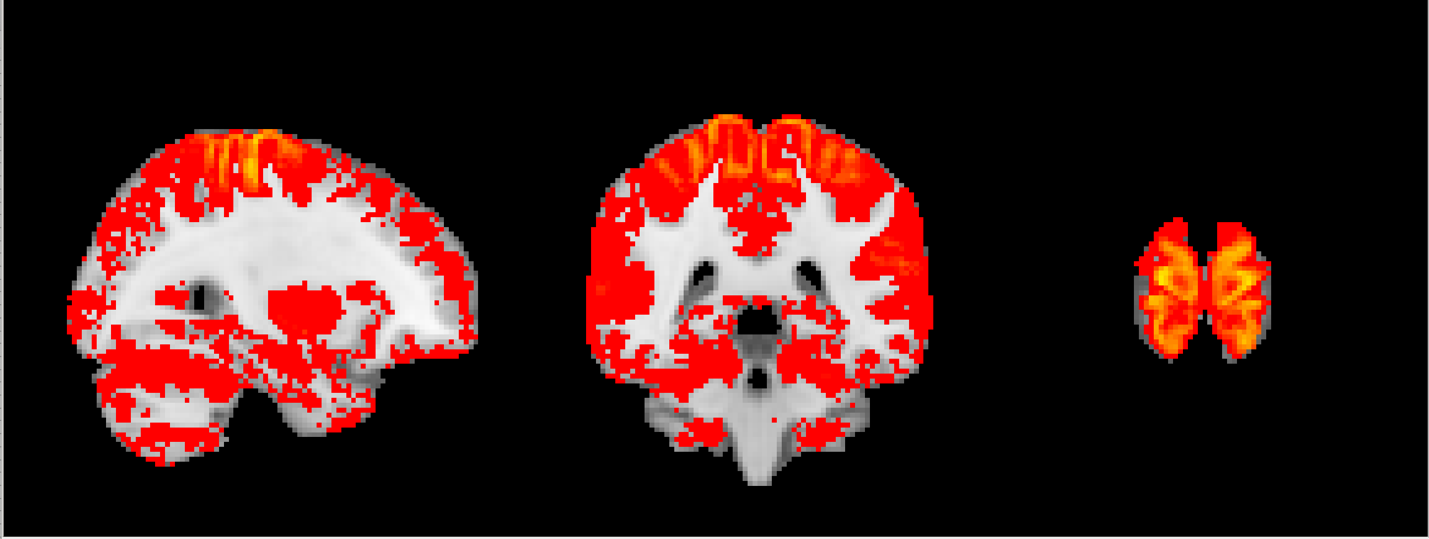

X= 30 mm

Y= -40 mm

Z= 76 mm

**VAN**

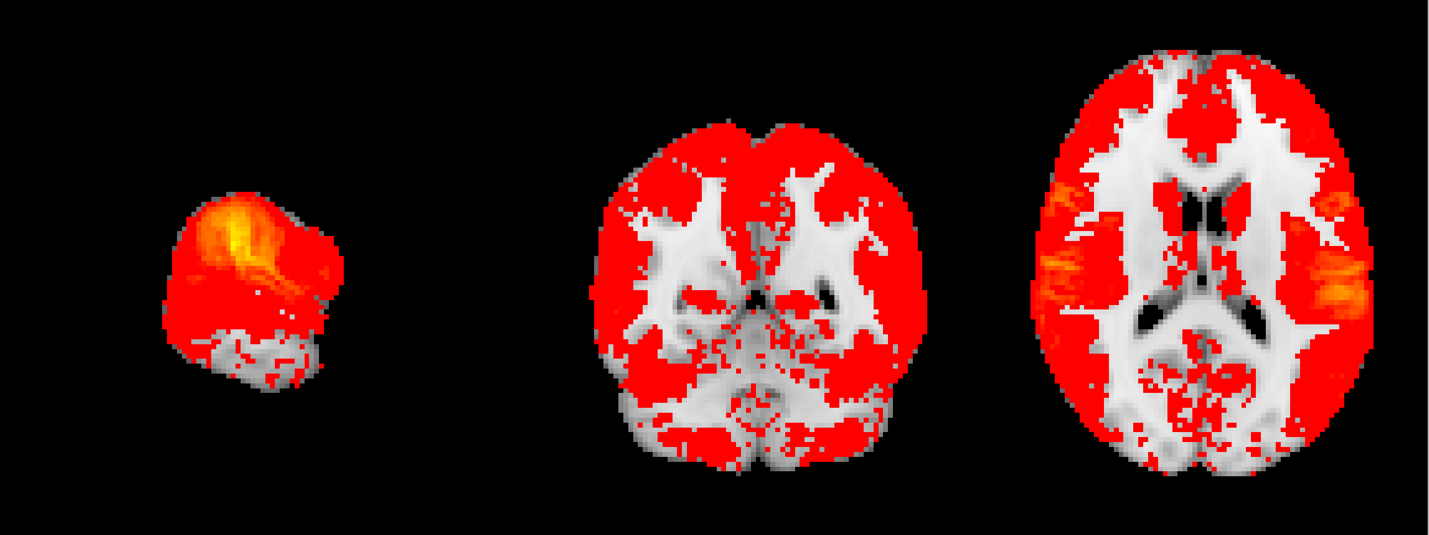

X= 63 mm

Y= -52 mm

Z= 17 mm

**VIS**

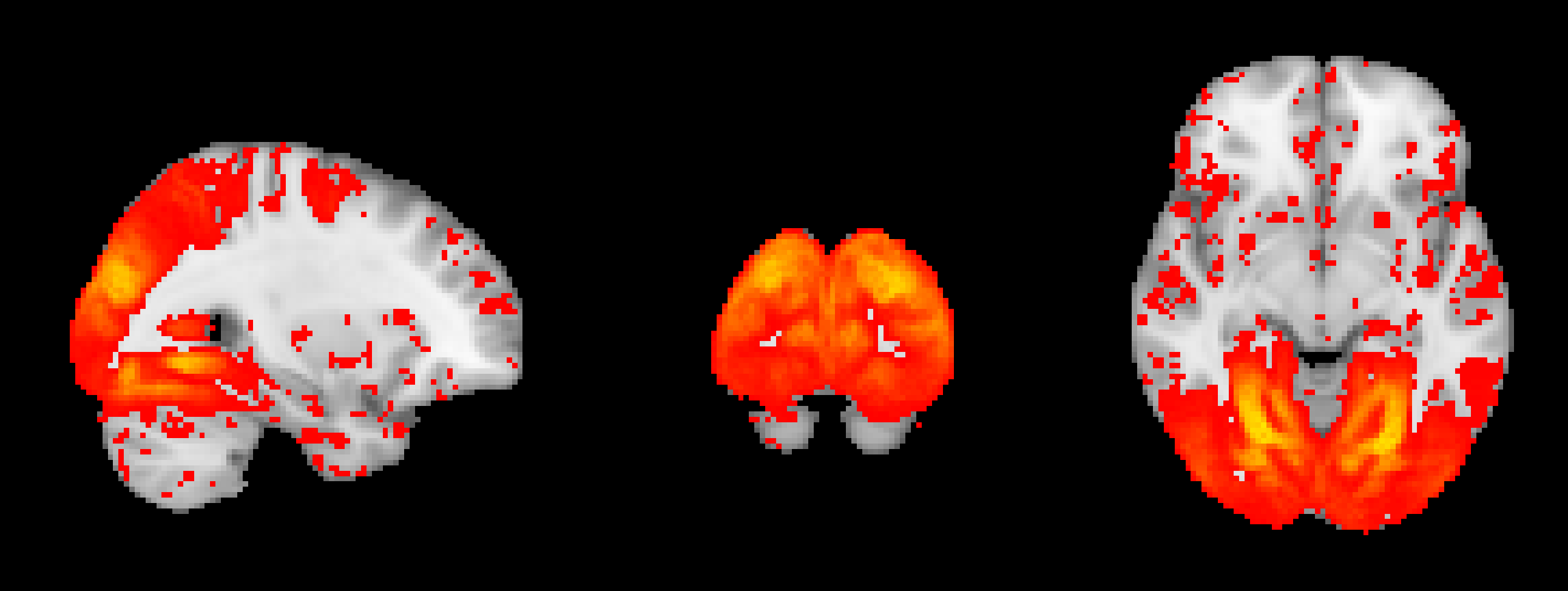

Z= -7 mm

X= 30 mm

Y= -90 mm

**Supplementary Figure S9:** Brain networks from Group ICA

**rCEN**

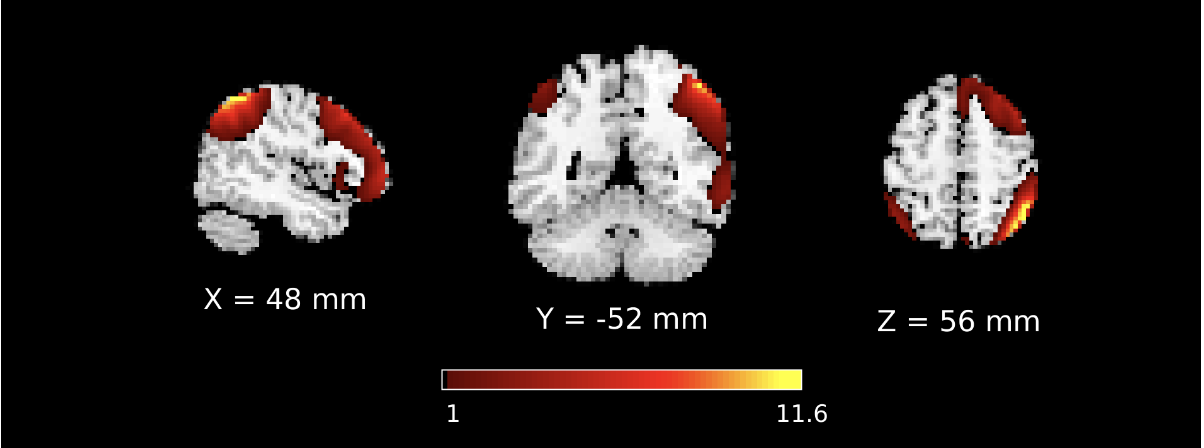

**lCEN**

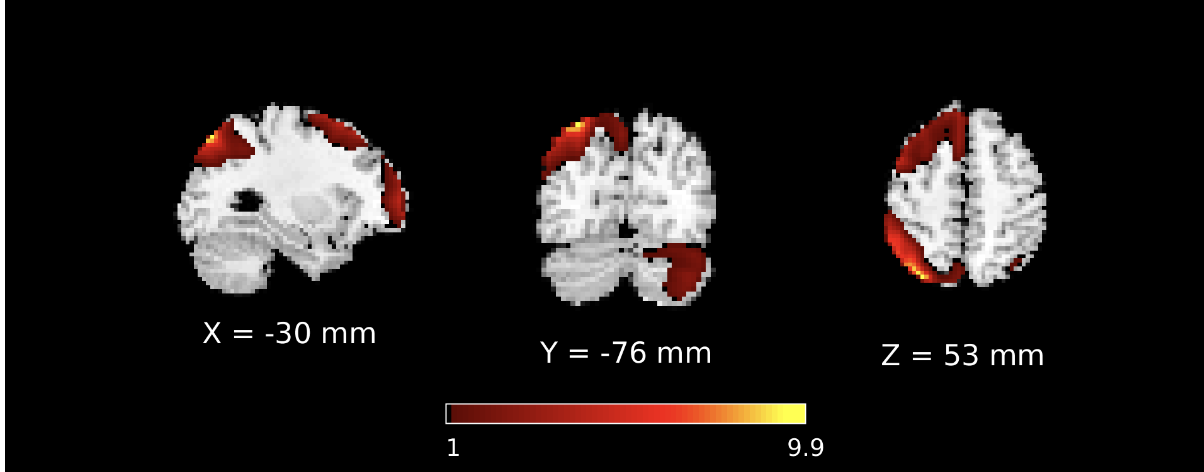

**DAN**

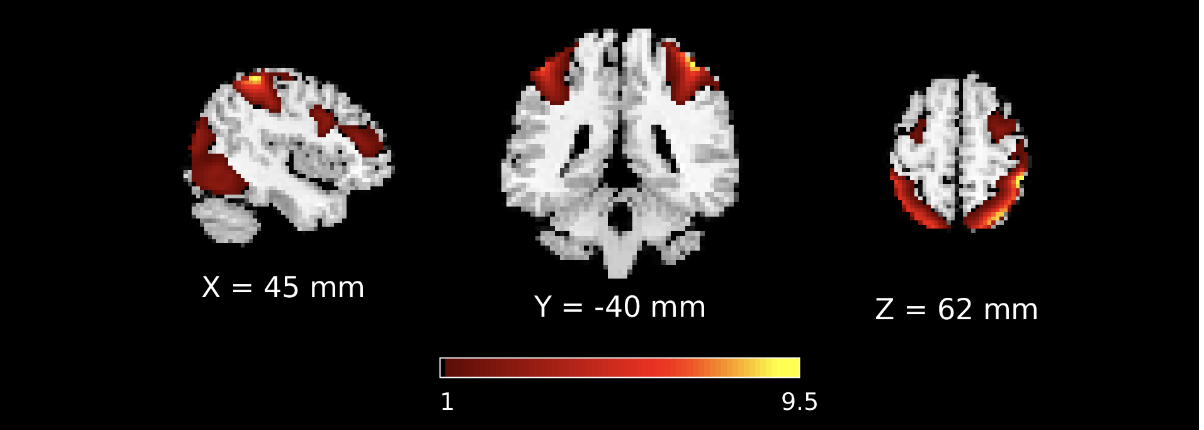

**aDMN**

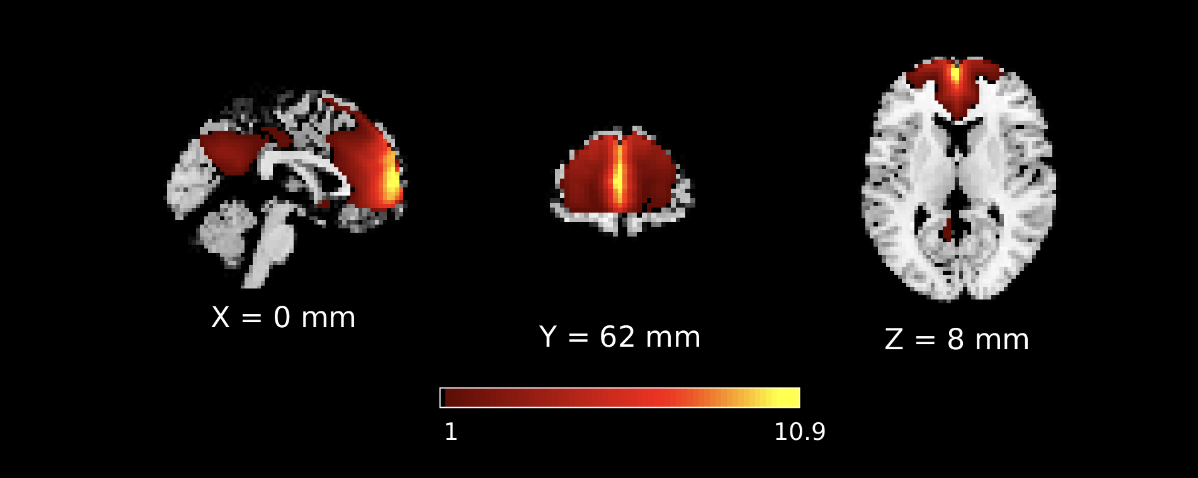

**pDMN**

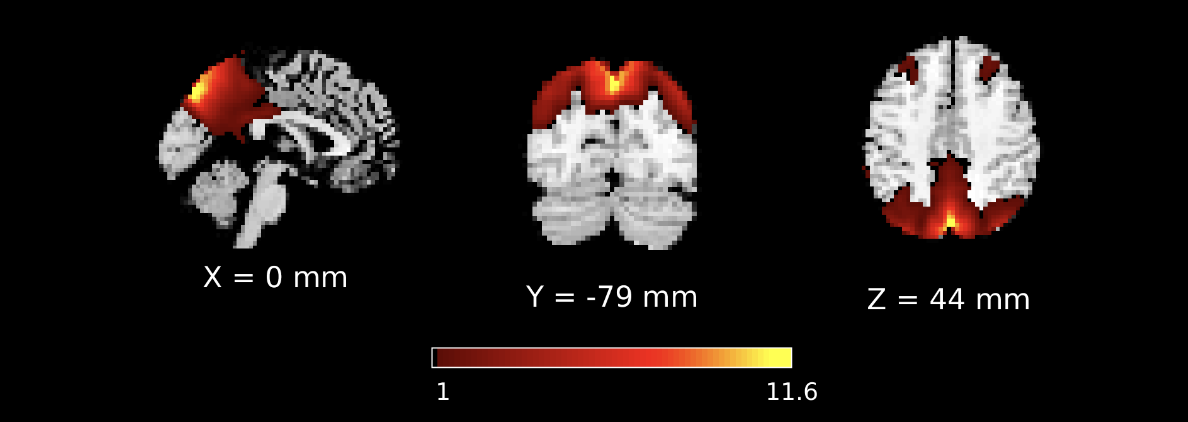

**SAL**

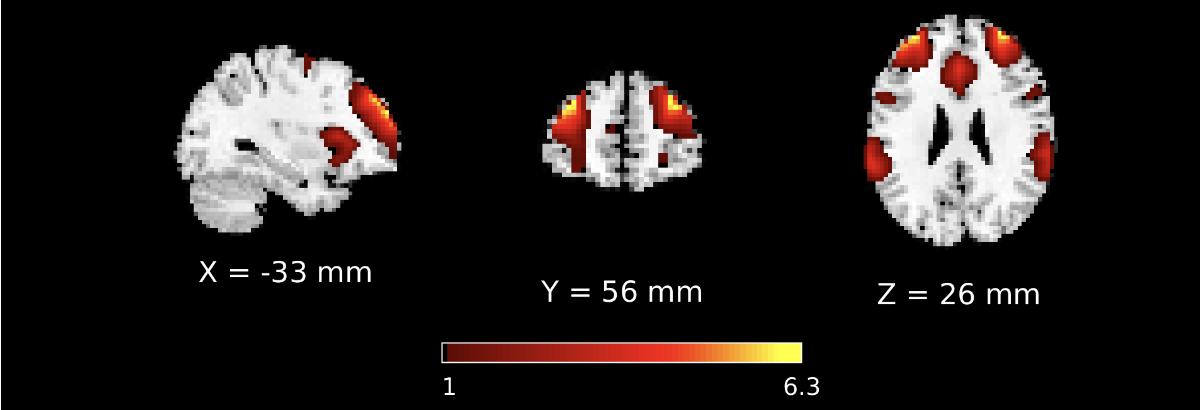

**SMC**

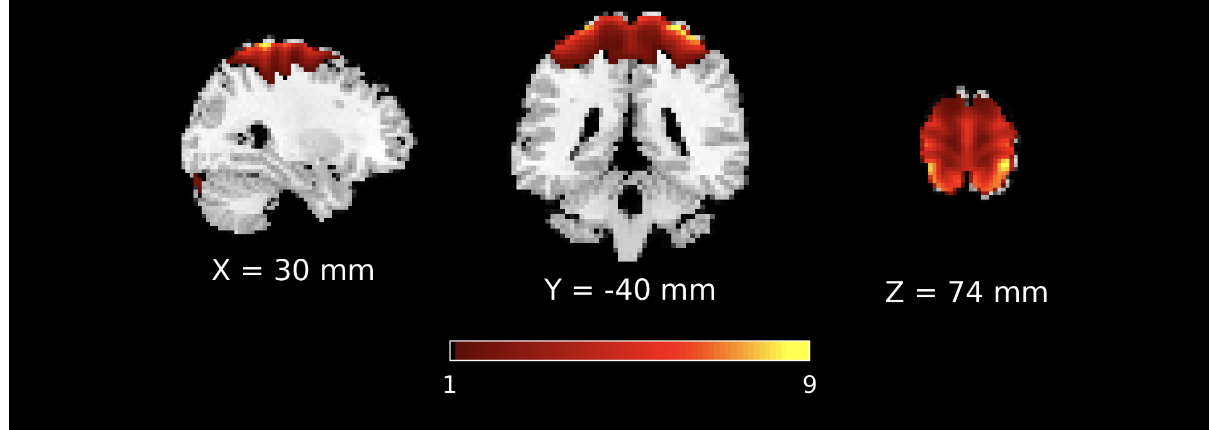

**VAN**

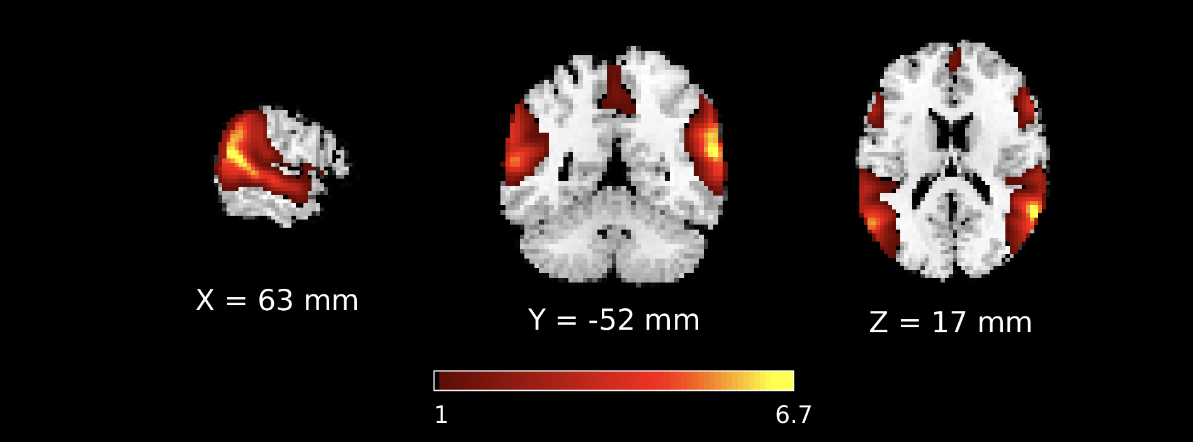

**VIS 1**

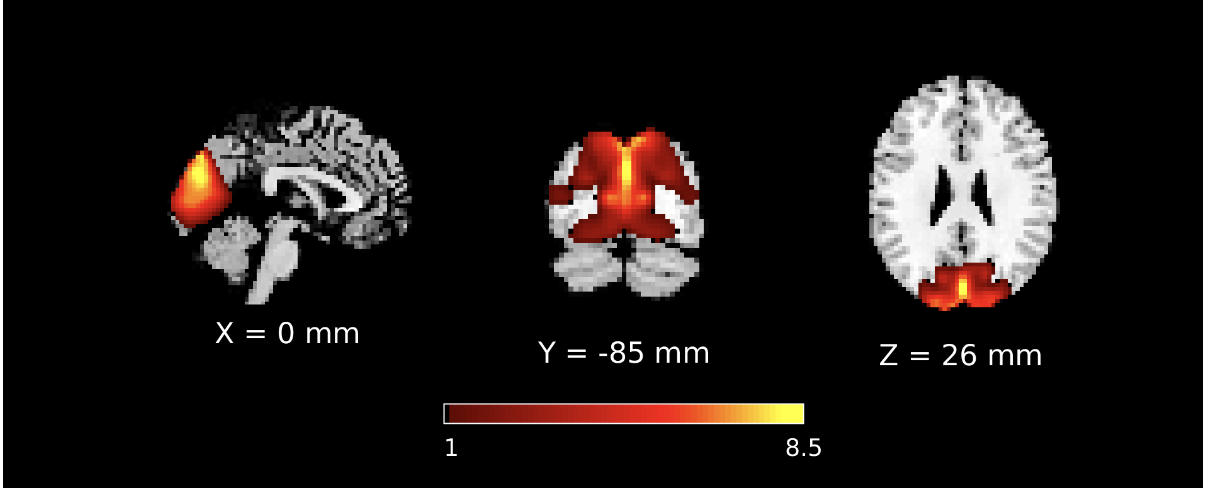

**VIS 3**

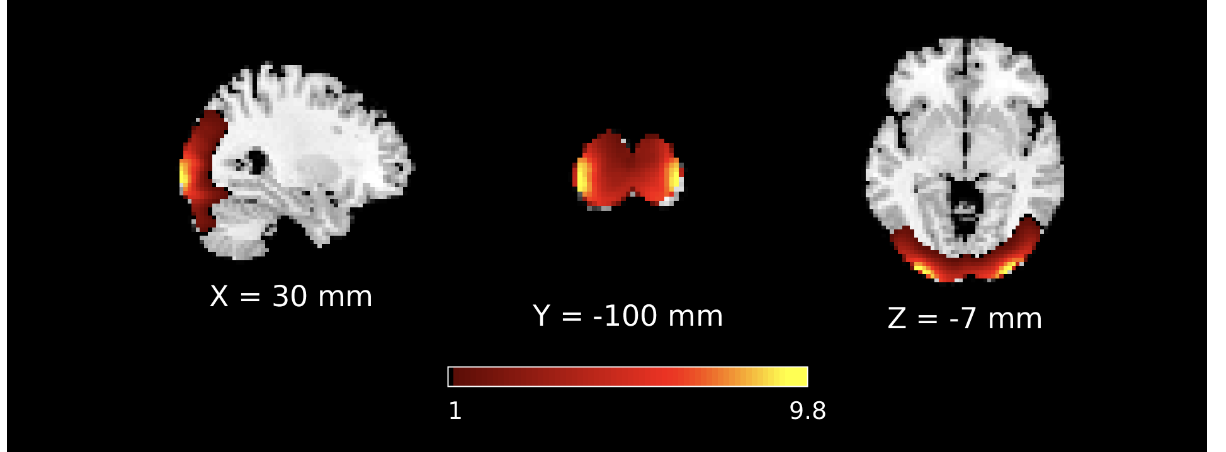
